# Tracking instances of task evoked oxygen influxes using a novel sensitive BLE enabled wearable fNIRS device identifies mPFC role in spatial memory

**DOI:** 10.1101/2025.11.27.690628

**Authors:** Rajesh Mandal, Gayathri Sai Prabhakaran, Arjun V Kowshik, Bhargav Makam Balaji, Balaji Jayaprakash

## Abstract

Real-world neuroimaging requires true portability and sensitivity to detect signals from neuronal activity. Conventional methods such as MRI, EEG, or PET are constrained by their size and instrumentation complexity. Near Infra-Red (NIR) based optical methods have the potential for sensitive detection with a smaller footprint. Despite rapid advances in NIR detection there is no known device that is implementable using off-the-shelf integrated microprocessors and possesses the above desired characteristics along with proven sensitivity to detect task evoked responses. Here, we present a Bluetooth Low Energy (BLE)-enabled, high-sensitivity wearable functional near-infrared spectroscopy (fNIRS) device designed for untethered cortical hemodynamic monitoring during naturalistic cognitive tasks. Our fully integrated optical sensing device merges optical data acquisition and wireless transmission into a compact, cable-free platform with a very small footprint. This enables continuous multi-channel recording without placing any constraint on the subject’s movement. We validate our device for reliable detection of task-evoked oxy- and deoxy-hemoglobin dynamics in the forearm, primary motor cortex, primary visual cortex, and prefrontal cortex. Subsequently, we capture real-time forebrain activity during a screen-based learning and memory task, revealing robust goal-specific hemodynamic responses. We formulate a method to identify the instances of peak neuronal activity and follow the “task Evoked Instances of Differential Oxygen influx(tEIDO)” as the subject is engaged in a task. These results highlight the potential of our proposed fNIRS device as a mobile neuroimaging solution for next-generation brain-computer interfaces and real-world cognitive monitoring.

**Graphical abstract:** 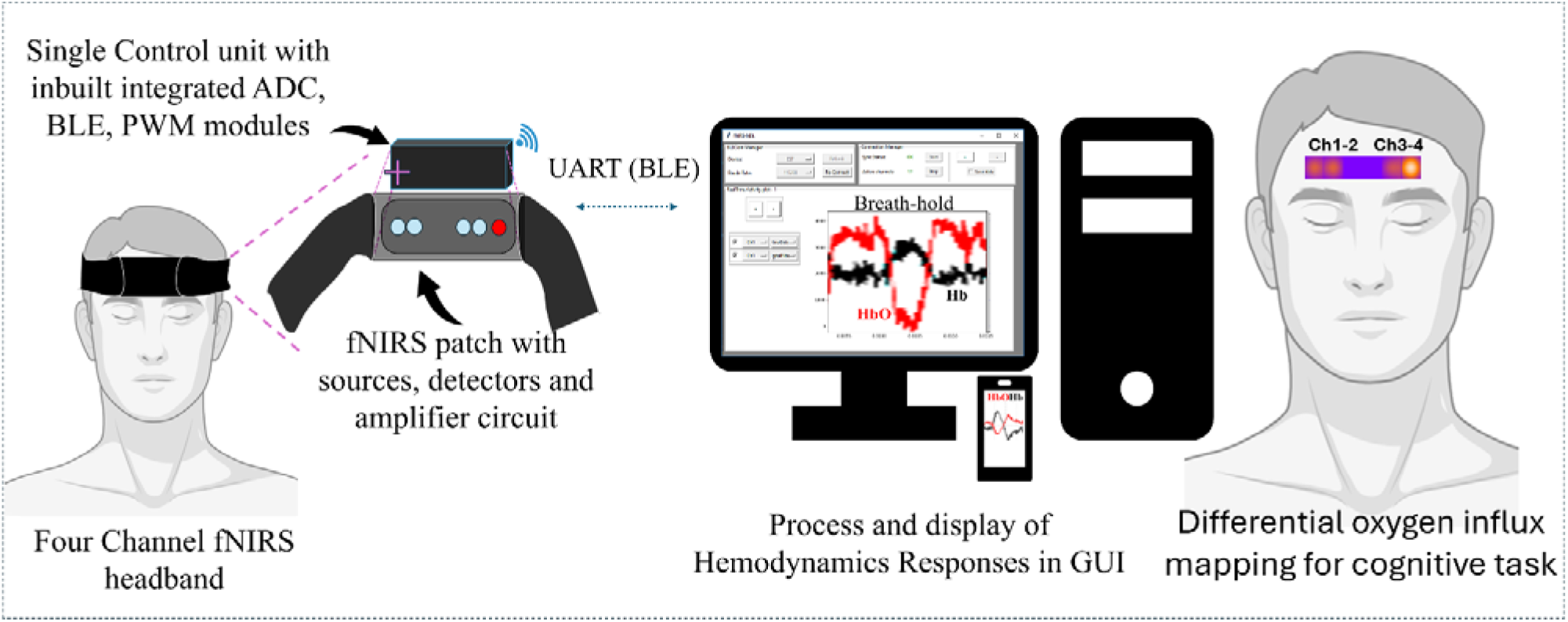

## 1. Introduction

Cognitive load-the mental effort required to process, store, and use information-directly shapes human performance^1^. In high-stakes settings, excessive demands can overwhelm working memory, slowing responses and increasing the likelihood of errors ^2^. Conventional neurocognitive assessments, including functional MRI (fMRI) and electroencephalography (EEG), have been foundational for studying workload but are limited to laboratory environments ^3,4^. fMRI, in particular, has played a central role in neuroimaging, yet its immobility renders it unsuitable for field applications^5^. This gap has driven the development of mobile neuroimaging techniques^6^ capable of capturing brain dynamics during naturalistic^7^ on-screen tasks. There exists a strong correlation between BOLD signal as measured in fMRI, Δ[HbO], Δ[Hb] and Δ[HbT] signals.^8^

Functional near-infrared spectroscopy (fNIRS) is a noninvasive optical method that measures cortical oxy- and deoxy-hemoglobin changes as proxies for neural activity^9,10^. Using near-infrared light, fNIRS enables continuous scalp-based monitoring of cerebral hemodynamics with good temporal resolution, without the constraints of huge scanner and restricted subject mobility^5,9^. While NIRS is inherently limited to shallow cortical regions (typically up to ∼3 cm), and the detected signal is influenced by all tissues and materials the light must traverse-such as scalp, skull, and even superficial factors like hair color, Owing to its strong correspondence with fMRI results, fNIRS is being used for ecologically valid monitoring of workload and fatigue ^2,5^. Being an optical technique the influence of absorbance properties of skin and hair color have been investigated and extent of their effects has been established^10^. Several optical ^11^and photoacoustic methods^12,13^ have been developed to monitor the blood flow, blood oxygenation state in model systems^2^.

Recent advances in wearable device design have transformed fNIRS into compact, wireless, and user-friendly systems^2,5,14,15^. But they still require one to integrate optodes, electronics, and wireless modules into lightweight head-mounted units. Portable devices have been shown to provide similar signal to noise ratio to table-top device^16^ but require relatively bulky setup and have not detected instantaneous task evoked instances of oxygen influxes from cognitive task. Further little is known about the sensitivity of these devices to detect neurological signals arising from cognitive tasks. These NIR devices either require tethering or backpacking of the associated electronics and are often power-intensive ^17^.

In cognitive neuroscience, fNIRS has been widely used to study workload, memory, and executive control^18,19^. The feasibility of monitoring brain activity during real-world cognitive tasks with a portable fNIRS platform was demonstrated^20^. However, the device remained relatively bulky, requiring participants to carry waist-mounted processing units.

A compact, bandage-sized Bluetooth-enabled fNIRS sensor was used^21^ for fatigue monitoring. However, the validation of this device was restricted to a single channel (single cortical region under observation), limiting its broader neuroimaging applicability. The maximum source detector separation of 20.2 mm, confines measurements to superficial cortical layers. In recent years, virtual environments - ranging from traditional on-screen setups to immersive virtual-reality platforms- have become powerful tools for studying cognitive processes in both humans and animal models ^22–28^. Capturing neuronal dynamics in real time while subjects interact with such environments provides a crucial window into the brain activity that underlies behavior. This integrated approach not only deepens our understanding of the neural mechanisms that support cognition but also establishes a foundation for diverse applications across experimental and clinical neuroscience. Thus, there is an unmet need to have a portable, compact, sensitive low power wireless device that can be built using readily available off the shelf integrated microprocessors for cognitive monitoring. The use of dedicated microcontrollers, ADC, source drivers and transmission modules increase the track length and thereby noise in the system. Thus, making it relatively less sensitive. Further the requirement to integrate these dedicated components limits the scalability as well as the ease of use thereby constraining the use to few laboratories. Here, we present a Bluetooth Low Energy (BLE)-enabled, high-sensitivity wearable functional near-infrared spectroscopy (fNIRS) device built using pre-integrated microprocessor. Our fully integrated optical sensing device merges data acquisition and wireless transmission into a compact, cable-free platform with a very small footprint. This enables continuous multi-channel recording without placing any constraint on the subject’s movement. The design can be used for untethered cortical hemodynamic monitoring during naturalistic cognitive tasks. Our device combines physiological, motor, and visual validations along with mapping of mPFC activity during a naturalistic cognitive paradigm.

We demonstrate the reliability of our device for detecting task-evoked oxy- and deoxy-hemoglobin dynamics in the forearm, primary motor cortex, primary visual cortex, and prefrontal cortex. This reliable detection validates the functionality of our device and capture real-time forebrain activity during a spatial memory task, revealing robust goal-specific instances of oxygen influxes in our hemodynamic responses during memory acquisition and retrieval. These results highlight the potential of our proposed fNIRS patch as a mobile neuroimaging solution for next-generation brain-computer interfaces and real-world cognitive monitoring.

## 2. Results

### Design and fabrication of the fNIRS headband

Our device design utilizes the off-the-shelf microcontroller ESP32 and its built-in ADC, BLE, and current-driving capacity of the DO terminals, along with a few other passive components, to orchestrate all aspects of fNIRS devices. The functional block diagram along with the real-world pictures of the fabricated head band is shown in Fig. 1. The functional block diagram (Fig. 1A) outlines the complete signal flow, starting from optical illumination through source diodes (S1 and S2), photon detection by silicon PIN photo detectors (Ch-#), and subsequent conditioning by the analog signal processing circuitry. Signals are digitized and processed in the onboard control unit before being wirelessly transmitted for storage and analysis. This architecture enables fully self-contained operation without reliance on external instrumentation. The device per se consists of two mechanically separable parts i) Front end – “the patch” and ii) Backend – “the body”. The patch consists of two sources, indicated by red rectangles, and four detectors, indicated by blue rectangles, all mounted behind a silicone mask, as shown in Fig. 1B. The detectors, Ch-1 and Ch-2, are placed at 0.5 and 1 cm (center-to-center distances) from the line joining the source centers. They correspond to 2 short channels, while the other two detectors, Ch-3 and Ch-4, are placed at 2.5 and 3 cm from the source center. These correspond to 2 long channels. The PCB, along with assembled components (ESP32, resistors, capacitors, diodes, and Op-Amps form the “body” of our device. This body is then packaged into a 3D-printed box. This box is then fixed to a Velcro band for ease of wearing and use. The whole device is then enclosed with an inner layer of Velcro band to enable adjustment of circumference while wearing, as shown in the front and rear views of the finished device in Fig. 1C. The user can recharge the device through the USB port available in the body. The Li-polymer battery for power is integrated within the body along with the control unit. Our design is modular, allowing for the straightforward replacement of individual components. Furthermore, this also enables future upgradation on the detector side, independent of the control unit, as they are interfaced through FFC cables and inter-joints. The fully furnished fNIRS biosensor weighs less than 80 grams, providing a portable and lightweight device for monitoring hemodynamics during cognitive task administration.

**Fig. 1.**
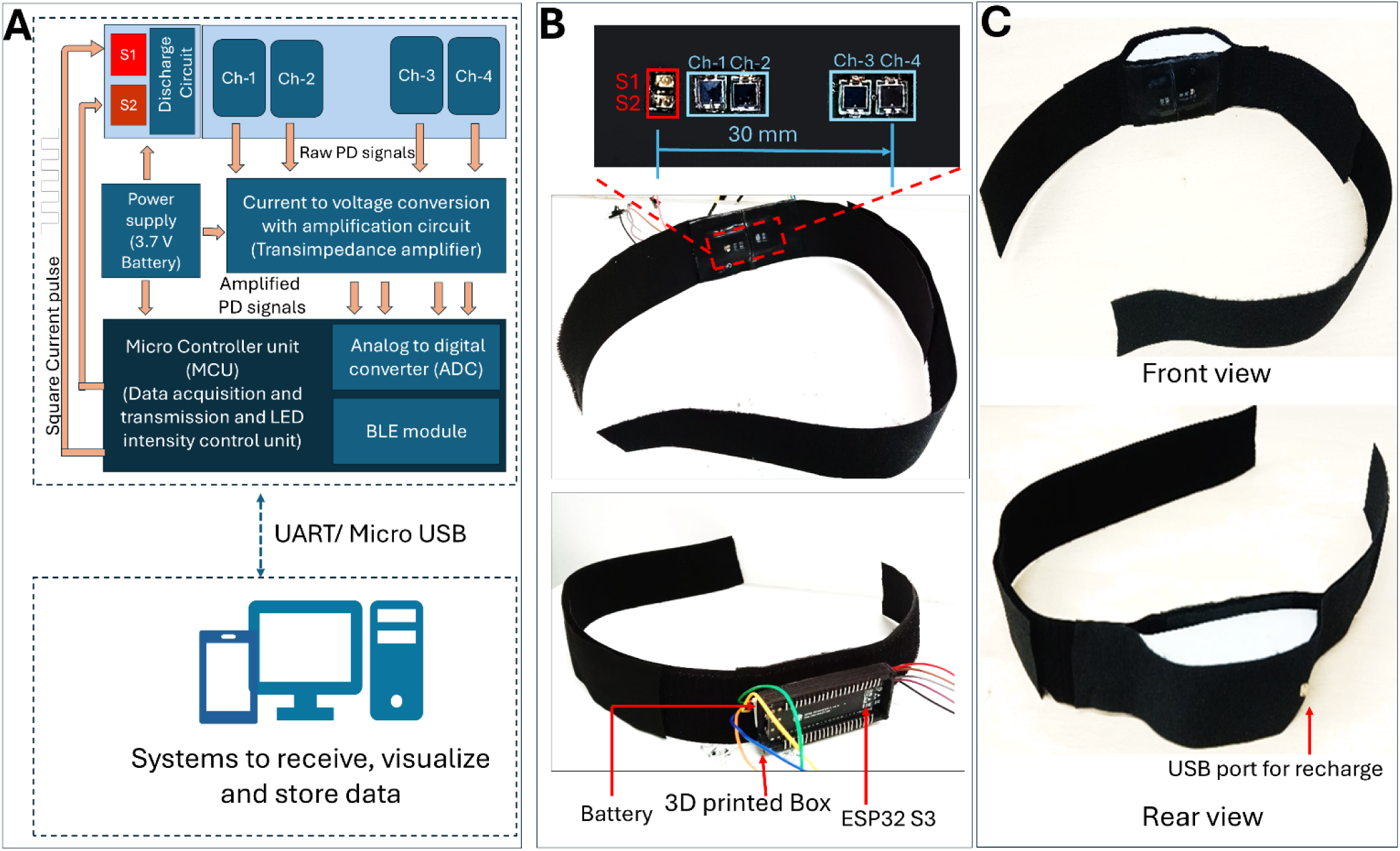
Overview and assembly of the fNIRS headband. (A) Functional block diagram illustrating each subsystem and the signal flow between components. (B) Assembly stages of the device: The upper panel displays the configured arrangement of optical sources and detectors, while the lower panel illustrates the integration of the control unit and battery. (C) Fully functional fNIRS headband: the top image reveals the internal scalp-contacting sensor array, and the bottom image displays the external housing.

### Validation of the developed fNIRS device and task-evoked hemodynamics

Next, we proceeded to validate our device using known interventions that alter either blood flow (occlusion) or oxygenation levels of the blood. For occlusion studies, we used the blood pressure monitor cuff to restrict blood flow during expansion, creating a transient and partial occlusion that is safe (Fig. 2A). For breath hold, participants were asked to hold their breath. We placed test, the headband on the participant’s forearm alongside a pulse oximeter module (MAX30102) available in the market (Fig. 2A). Once the trial begins the cuff was pressurized using a motorized BP monitor and following the release of the cuff pressure (after 300 s from the start of the experiment) the participant was instructed to hold their breath (for 30 s). Representative time courses obtained from four fNIRS channels during this period are shown in Fig. 2B. Two small regions from Ch-4 marked in green and purple rectangles corresponding to occlusion and breath-hold instances are zoomed out and presented at the bottom. We observed that cuff inflation (indicated by the vertical dashed lines in Fig. 2B) elicited a clear and complex hemodynamic response, with deoxyhemoglobin (ΔHb) (blue line) increasing by +60±10.1 nmol/L measured across (N = 5, trials/participants) and hemoglobin (ΔHbO) decreasing by −36.48±6.2 nmol/L post occlusion. Simultaneously, we measured arterial oxygen saturation (SpO₂) using a commercial pulse oximeter. The saturation dropped by ∼5% (from a baseline of 99.83% to 94.4%). We note that this is consistent with our fNIRS-derived hemodynamic changes. The complex dynamics associated with oxygenation/deoxygenation are absent, illustrating the higher temporal resolution and faster response offered by our headband. Our analysis of the fNIRS data during breath hold (300 - 330 s) showed a clear hemodynamic response (Fig. 2B bottom right panel-zoomed out purple rectangle), with deoxyhemoglobin (ΔHb) concentration increasing by 12.6±5.6 nM and concentration of oxyhemoglobin (ΔHbO) decreasing by 13.4±0.9 nM. The arterial oxygen saturation (SpO₂) measured simultaneously by the pulse oximeter dropped by ∼3% (from baseline 99.83% to 96.8%), consistent with the fNIRS-derived hemodynamic changes.

**Fig. 2.**
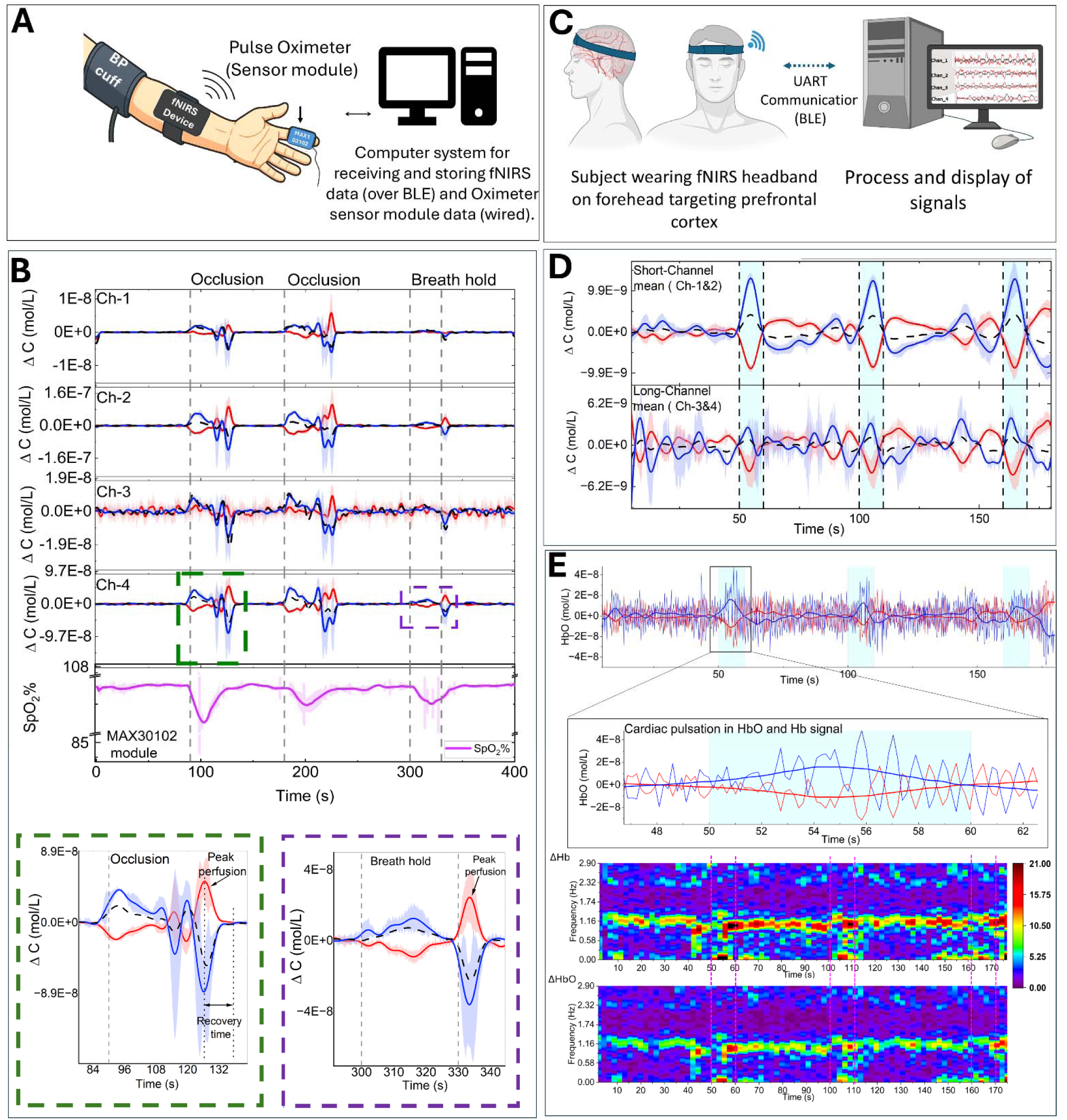
Validation of the developed fNIRS biosensor and task-evoked hemodynamics. (A) Experimental setup for vascular-occlusion validation: fNIRS headband, pulse-oximeter probe, and blood-pressure cuff co-located on the participant’s forearm. (B) Time courses (mean ± SEM across trials) of hemoglobin changes from four fNIRS channels (Ch-1 to Ch-4): Δ[HbO](red), Δ[Hb](blue), and Δ[HbT] (black, dashed). The bottom trace shows pulse-oximeter arterial oxygen saturation (SpO, magenta). The vertical dashed line marks cuff inflation (onset of occlusion) and the breath-hold event. An occlusion and breath-hold response from Ch-4 is highlighted at the bottom. (C) Breath-hold experiment: fNIRS headband on the forehead with real-time streaming to a PC. (D) Breath-hold responses (mean ± SEM across trials): concentration changes ΔHbO, ΔHb, and Δ[HbT] from the short-channel (top) and long-channel (bottom). Blue-shaded windows between vertical dashed lines denote 10-s breath-hold periods. (E) Cardiac pulsation in the raw fNIRS signal (top) and the smoothed trace after pulsatility suppression; corresponding time frequency spectra (bottom) show a dominant peak centered at ∼1.2-1.4 Hz consistent with heart rate. 2 male subjects; Subject 1 (26 years) completed five trials of the vascular-occlusion validation, and Subject 2 (28 years) completed five trials of the breath-hold experiment.

It was observed that immediately after the cuff pressure was released or breath release, all fNIRS channels detected a marked rebound in blood flow, characterized by an overshoot in ΔHbO and a corresponding decrease in ΔHb. Interestingly, this post-occlusion response was not captured in the pulse oximeter SpO₂% measurements. A single trial trace from any of the channels highlights the robustness of these responses to both occlusion and breath-hold events, showing peak perfusion and recovery time.

After the above measurements, we assessed the device sensitivity for measuring cortical events. For this, we positioned the headband on the forehead during a controlled breath-hold task (Fig. 2C). Average concentration changes measured over 5 trials are shown in Fig. 2D. During 10-second breath-hold periods (blue-shaded regions), we observed a decrease in oxy-hemoglobin (ΔHbO) followed by an increase in hemoglobin (ΔHb) in all four channels.

The raw ΔHbO and ΔHb traces displayed periodic high-frequency physiological oscillations. Analyzing the data further, we attribute these to cardiac pulsation, which can also be resolved (Fig. 2E). These physiological fluctuations could be suppressed following a temporal smoothing (adjacent averaging, 26 points (corresponding to ∼0.2 Hz)), providing high SNR hemodynamic traces. The time-frequency spectrum of the raw signal from one of the subjects is shown in Fig. 2E. The spectrum (SFig. 4) exhibits a dominant band centered at ∼1.3 Hz in both ΔHb and Δ[HbO] time series measurements, corresponding to a heart rate of ∼78 beats per minute. We note that although the intensity of these bands from ΔHb and Δ[HbO] are different, the time evolution and the center frequency are very similar. Thus, with our device, we could capture slow hemodynamic responses as well as detect fast vascular dynamics.

### Motor Cortex Activation During Finger Tapping

We next proceeded to validate responses resulting from distinct neural activity. We utilized a well-established rhythmic finger-tapping task that is known to specifically engage the motor cortex ^29^. Previously, 2 Hz was identified as the optimal frequency for focal activation of primary motor cortex (BA 4) during rhythmic finger tapping. We implemented a block-design finger-tapping task tailored for fNIRS. Briefly, the task comprises five sub-blocks of 24 tap cycles totaling 120 tap cycles spread across the entire task. Each session consisted of 24 cycles of 12-second activation (tapping) followed by 12-second rest (hands flat), for a total run time of 9.6 minutes. Fig. 3 summarizes the hemodynamic responses recorded over the primary motor cortex during one such block in one of the participants.

**Fig. 3.**
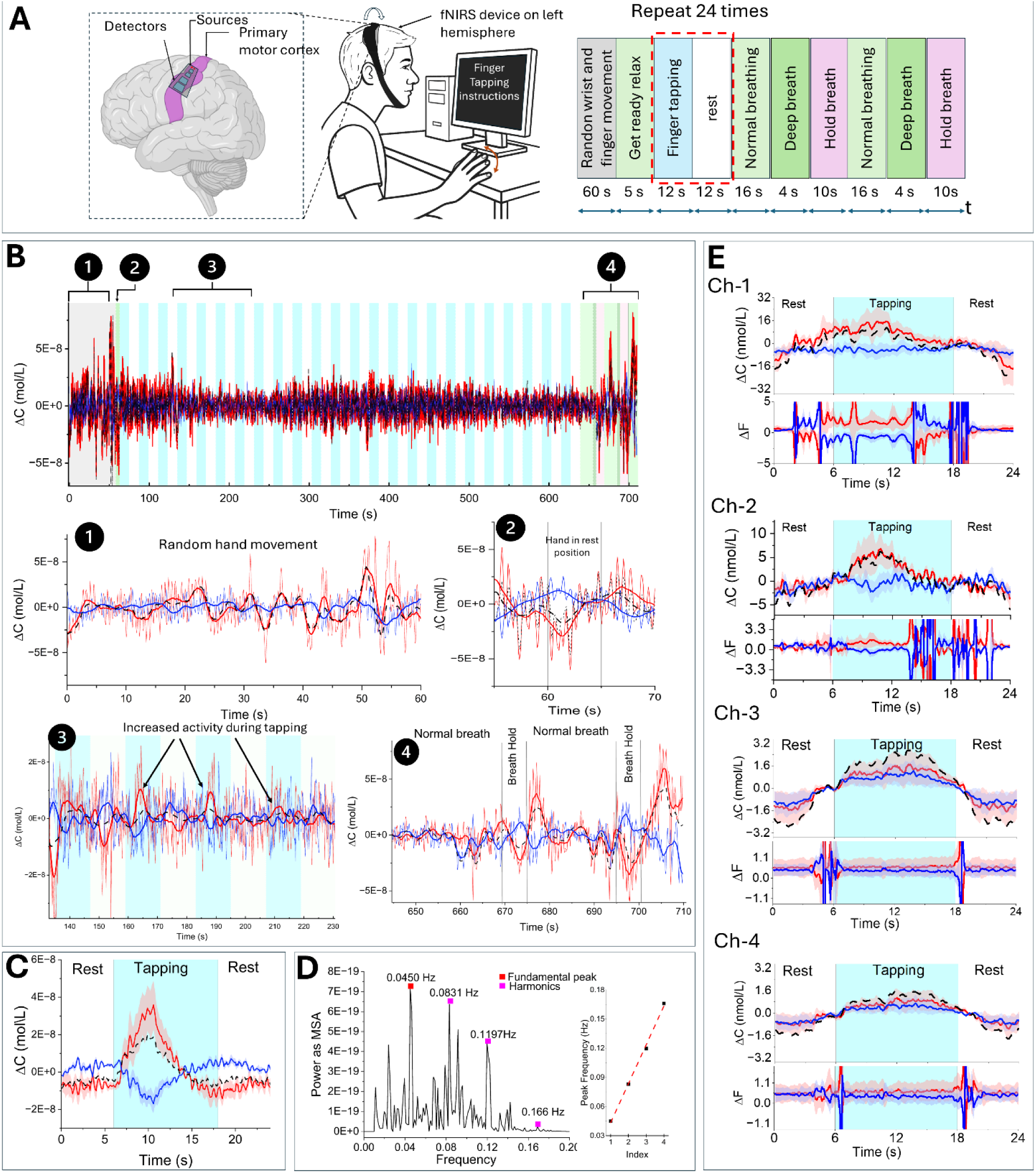
Hemodynamic responses in the primary motor cortex during finger tapping. (A) Setup: participant wears an fNIRS headband over the primary motor cortex and performs rhythmic finger taps guided by on-screen cues (timing protocol shown at right). (B) Representative trace from Ch-2: Δ[HbO](red), Δ[Hb](blue), and Δ[HbT] (black, dashed). Events 1-4 are highlighted and shown as short segments; bold lines represent the smoothed versions of the raw signals. (C) Epoch-averaged response across 24 tap-rest blocks (mean ± SEM across epochs) of (B). (D) Power spectrum of Δ[HbO]showing the task fundamental at ∼ 0.042 Hz (1/24 s) and its harmonics. (E) Across participants, the mean ± SEM traces are shown for Δ[HbO](red), Δ[Hb](blue), and Δ[HbT] (black, dashed). The panels below display the corresponding mean ± SEM of Δ[HbO](red) and Δ[Hb](blue) after normalization by Δ[HbT] for channels Ch-1 through Ch-4. Vertical dashed lines mark alternating rest and tapping periods. (n=14 participants,8 male, 6 female, age= 24-29 years).

We positioned a four-channel fNIRS patch over the precentral gyrus-targeting BA 4-using 10-20 EEG landmarks (C3/C4) (Fig.3A). Towards the end of the task, the subjects were asked to do a breath hold for 10s twice with a gap of 20 sec as a positive control to ensure the device is coupled to scalp efficiently.

The time evolution of Δ[Hb] and Δ[HbO], as measured from Channel 2, is shown in Fig. 3B. We demarcate the response into distinct temporal segments (denoted by the numeric labels in Fig. 3B) for closer and detailed examination. These segments correspond to different phases of the task. Bold lines represent smoothed traces, which clearly delineate task-related hemodynamic modulation from baseline fluctuations. In all segments, we observed clear and distinct task-evoked dynamics that were specific to the phase of the task. We observe a specific increase in oxyhemoglobin (ΔHbO, red) and a decrease in deoxyhemoglobin (ΔHb, blue) during most of the tapping blocks (blue regions in Fig. 3B, segment-3). We also observe an increase in deoxyhemoglobin and a decrease in oxyhemoglobin during rest blocks (white regions in Fig. 3B, segment 3). Segment 4 shows the change in ΔHb and Δ[HbO] concentrations during the breath hold. Next, we averaged responses across 24 tap-rest blocks (Fig. 3C). The average clearly demonstrated the expected neurovascular coupling signature, characterized by increased oxygen delivery and reduced deoxygenation during motor activity.

The power spectrum of ΔHbO (Fig. 3D) revealed a dominant frequency at 0.045 Hz (close to 0.42 Hz), corresponding to the 24-second tap-rest cycle, along with its harmonics. Across-subject averages (n = 13 participants) yielded consistent results, showing a ΔHbO increase in all channels. Vertical dashed lines in Fig.3E mark the tap and rest blocks, aligning well with the cyclical hemodynamic responses.

### Primary Visual Cortex Activation During Dynamics-Contrast Stimuli

Fig.4 shows the hemodynamic responses recorded from the primary visual cortex during a dynamic-contrast visual stimulation task. We drew on previous fMRI investigations of visual cortex organization-^30^ ^31^ for high-resolution retinotopy in V1. For network dynamics during feature binding, both of which target Brodmann areas 17-19.

**Fig. 4.**
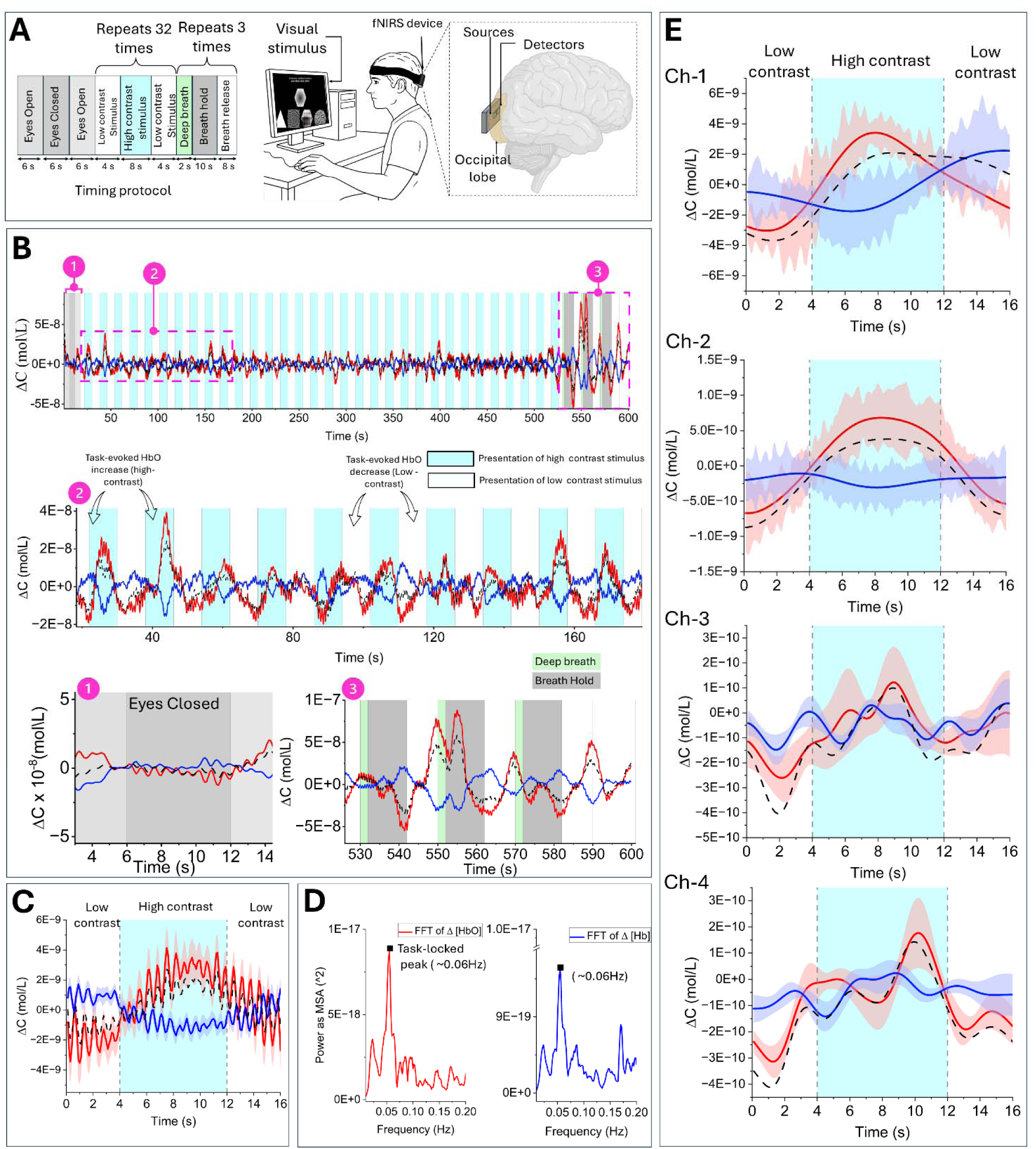
Hemodynamic activity in the primary visual cortex during a dynamic-contrast visual task. (A) Experimental setup: the participant wears the fNIRS headband over the occipital region and views stimuli on a monitor. (B) Representative trace from Ch-2: Δ[HbO](red), Δ[Hb](blue), and Δ[HbT] (black, dashed). Events 1-4 are highlighted and shown as short segments. (C) Epoch-averaged response across 32 low contrast-high contrast-low contrast blocks (mean ± SEM across epochs) of (B). (D) Power spectrum of Δ[HbO]and Δ[Hb]showing the task fundamental at ∼ 0.06 Hz (1/16 s). (E) Across-subject averages from four channels (Ch-1, Ch-2, Ch-3, and Ch-4): Δ[HbO](red), Δ[Hb](blue), and Δ[HbT] (black, dashed). Vertical dashed lines mark alternating rest and tapping periods. (n=16 participants,9 male, 7 female, age= 24-29 years).

The fNIRS headband was positioned over the primary visual cortex while participants viewed alternating blocks of low- and high-contrast images on a monitor (Fig. 4A). Each block lasted 16 seconds, with 32 low-high-low contrast transitions per epoch.

A typical time course from Channel 2 is displayed in Fig. 4B. In response to high-contrast visual stimulation, the concentration of oxyhemoglobin (ΔHbO, red) increased, while that of deoxyhemoglobin (ΔHb, blue) decreased. We also observe a rise in total hemoglobin (Δ[HbT], black dashed), reflecting an enhanced cortical blood volume. Highlighted segments (Events 1-4) illustrate repeated, well-defined responses across consecutive stimulus presentations.

When averaged across 32 contrast blocks, the visual-evoked response exhibited a robust increase in oxyhemoglobin (Fig. 4C). These findings are consistent with the expected neurovascular response in the primary visual cortex.

Spectral analysis of the ΔHbO trace revealed a dominant peak at 0.06 Hz, corresponding to the 16-second block cycle (Fig. 4D). Group-level analysis across seventeen participants confirmed reliable activation across all four channels (Fig. 4E).

### Task-evoked Instances of Differential Oxygen Influx (tEIDO)

Encouraged by the clear hemodynamic response that exhibits clear decoupling between oxy and deoxy hemoglobin responses in individual subject responses (Fig. 5A), we proceeded to identify the instances of peak neuronal activity. Previously, neuronal activity has been shown to trigger an increased inflow of oxygenated blood in local regions that are modelled as a Balloon^32^. This results in a characteristic time dependence of the decoupling between oxy and deoxy hemoglobin ^33^. Here, we make use of this decoupling and our ability to measure (Eq. 8) it distinctly during a task to identify the instances of peak neuronal activity.

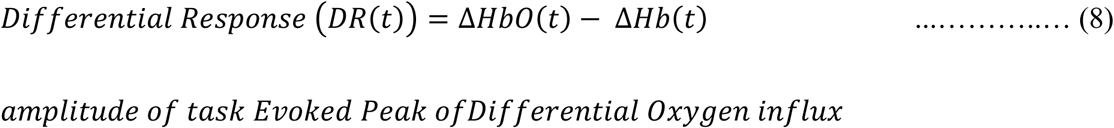

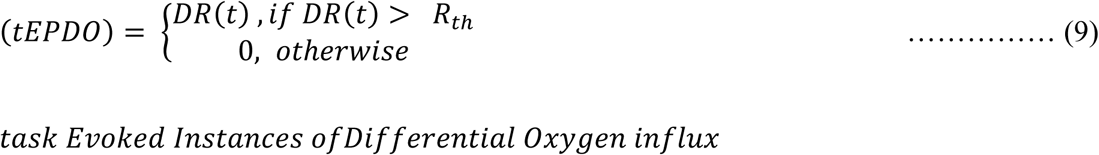

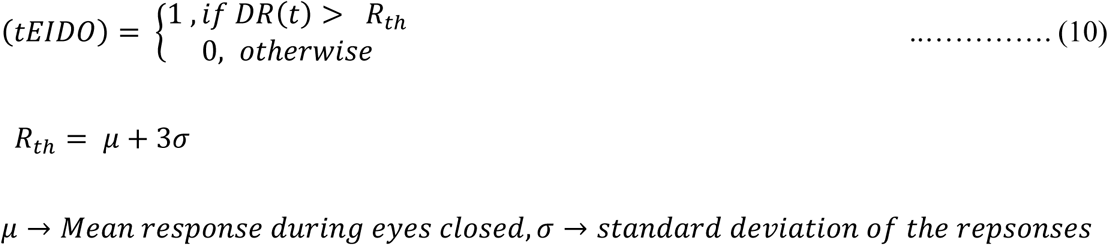

**Fig. 5.**
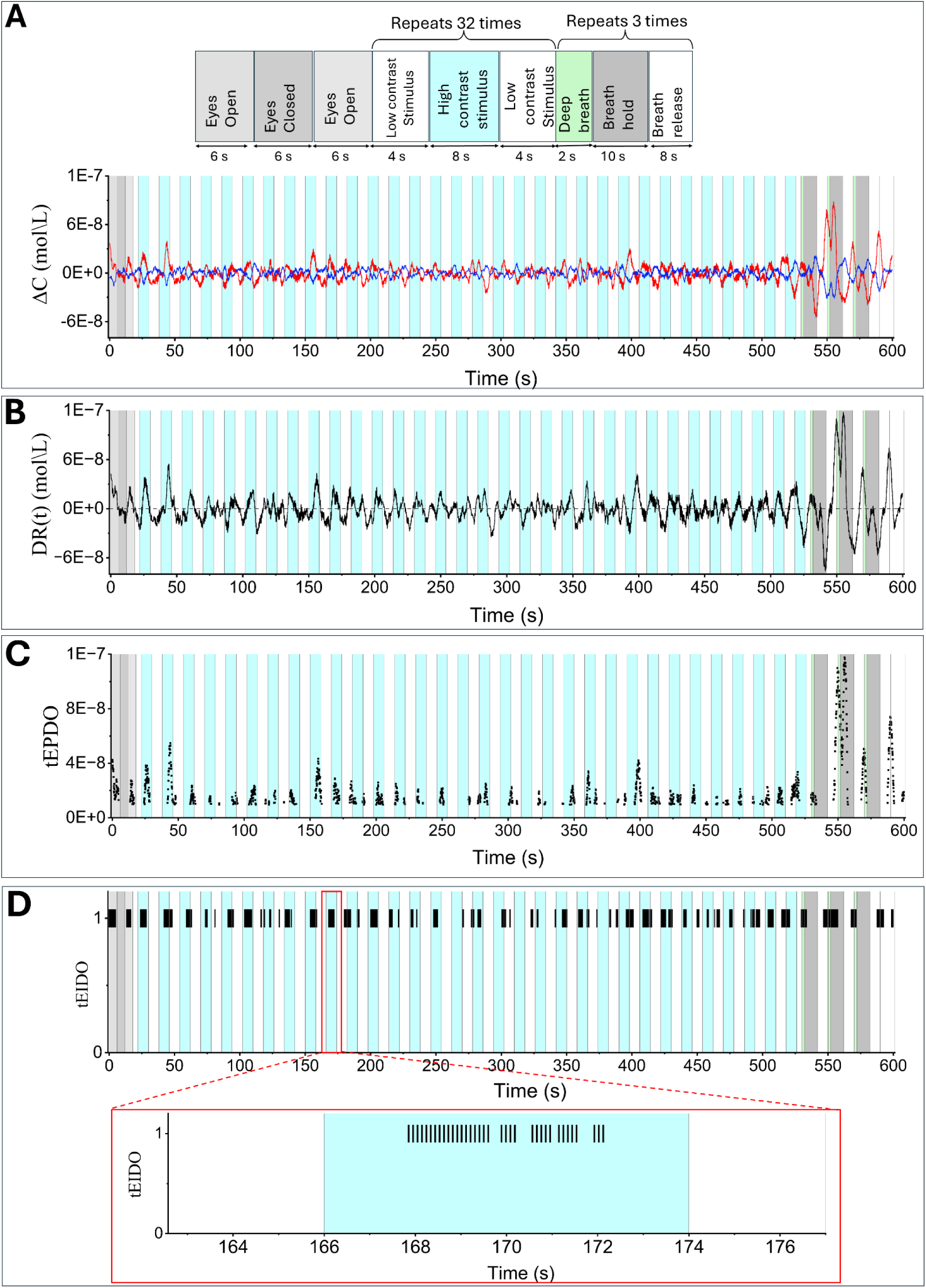
Task-evoked Instances of Differential Oxygen Influx (tEIDO). (A) Single-channel trace (Ch-2): Δ[HbO](red) and Δ[Hb](blue) during the task. (B) Differential response (DR(t) = ΔHbO(t) - ΔHb(t)). (C) Detected task-evoked Peaks of Differential oxygen influx (tEPDO) using a fixed threshold at μ_baseline_ +3 x a_baseline_ (baseline: eyes-closed state). (D) task-evoked Instances of Differential Oxygen Influx (tEIDO).

In Fig. 5B, we plot this differential response measured during the dynamic contrast visual task described above for a single subject. We use the responses obtained during the eye-closed period and measure the mean (μ) and the standard deviation (σ) of these differential responses during this period. Subsequently, we use them to determine the threshold (*Rth*) to classify the responses as tEPDO (Fig. 5C) as per Eq. 9. We identify the periods /instances at which tEPDO occurred as the task-evoked periods/instances of higher neuronal activity(tEIDOs) as per Eq. 10. These are shown in Fig. 5D. We note that tEIDOs could represent either a transient isolated increase in oxygen influx or could be due to a sustained increase, both of which result from an underlying neuronal activity^32,34^.

### Location-specific activation of cortical sub-regions in a screen-based navigational task

Next, we probed the involvement of the prefrontal cortex (BA 10) during a spatial navigation task. The prefrontal cortex is known to be involved in facilitating the retrieval of spatial information and in reaching the destination. We developed and utilized a virtual maze task that is similar to the Morris Water Maze task, which is extensively used in rodents as a test for memory ^35^. The on-screen task is described in the methods, but briefly, it consists of four trials in one session. The subject learns to navigate in virtual ground to locate a hidden treasure using distal cues (illustration in Fig. 6A). The first 3 trials (T0 – T2) constitute the training phase (Fig. 6A schematic) while the fourth trial (T3) is the probe/testing trial. During all these trials, we placed our device on the subject’s forehead while they were engaged in the task. We see the subjects learn to navigate to the treasure as the trial progresses. We follow this progress by measuring the residence time heat map across the trials. The average residence time heatmaps (Fig 6B) show the emergence of selective search. We quantify this by comparing the average time spent searching by the subject in each quadrant. When we compared this across the quadrants at the probe trial, we observed that Q2 residence time was significantly higher than all other quadrants (*one-way ANOVA, F*(3, 56) = 55.90, *p* < 0.0001; post hoc Tukey). Moreover, Q2 was also significantly above chance (65.68 %) (*t*-test, *p* < 0.0001, *n* = 15). Thus, we observe that the subjects have learned to navigate and reach the treasure location. During the probe or testing trial (T3), a marked reduction in proximity to the treasure location was observed, indicating a clear consolidation of spatial memory. One-way ANOVA revealed a significant effect of trial on proximity (*F*(3, 62) = 10.51, *p* < 0.0001).

**Fig. 6.**
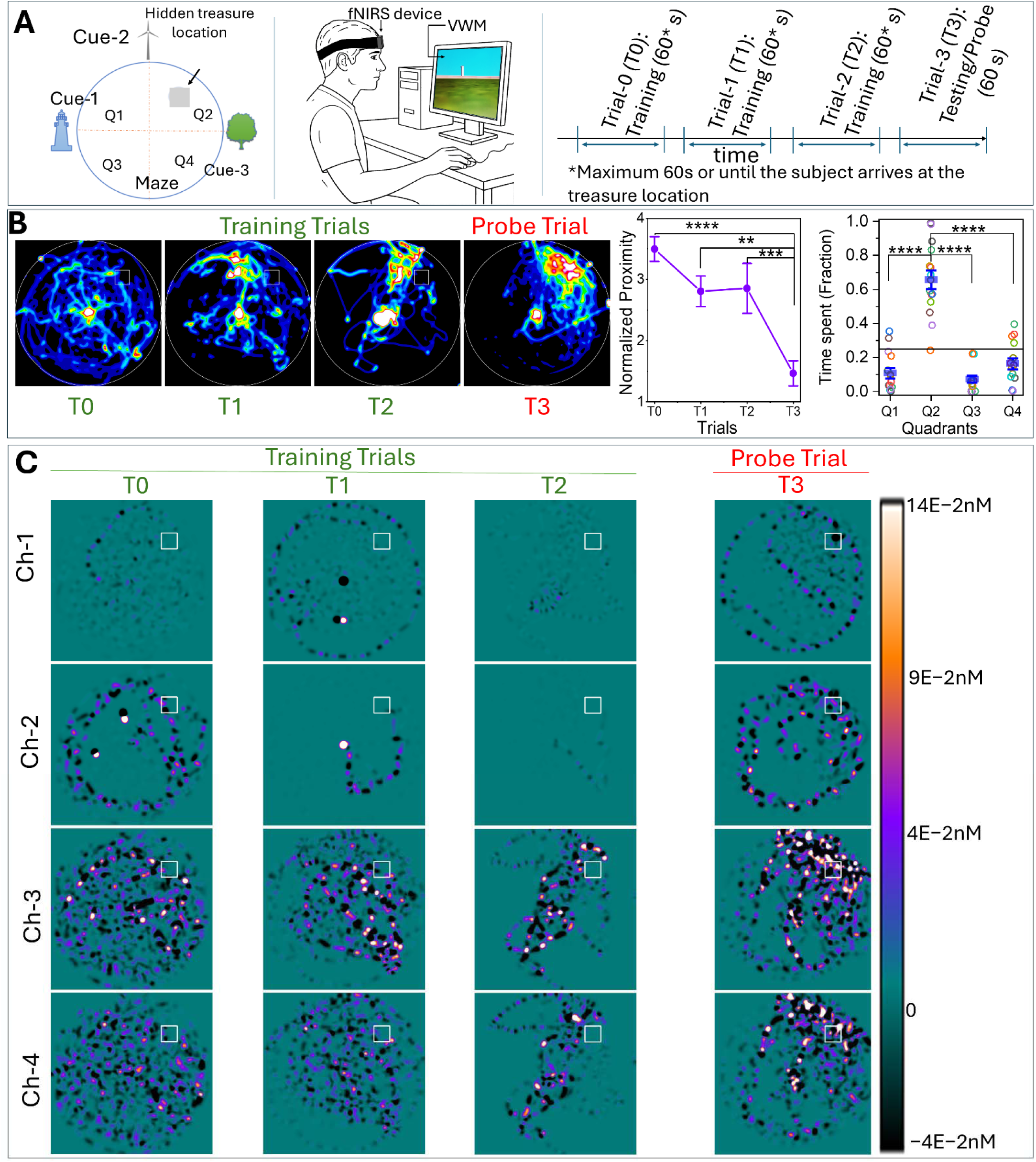
Location-specific activation of cortical sub-regions in a screen-based navigational task. (A) Experimental setup: Left: Schematic of the open-field virtual maze with a hidden platform and distal visual cues. Centre: Subject performing the task on a computer while wearing the fNIRS headband over the frontal cortex. Right: Timing protocol for a single session. (B) Mean residence heatmaps for Trials 0-3 (left to right), illustrating the subject’s spatial occupancy over the maze during each trial (left). During the probe/testing trial, the center of mass of these maps shifts significantly closer to the treasure location, with normalized proximity showing robust reductions (***p < 0.0001, ***p < 0.001, **p < 0.01; middle). Consistently, the proportion of time spent in Quadrant 4 (Q4) rises significantly, exhibiting a substantial increase (****p < 0.0001; right). (C) Residence time normalized differential oxygenation (DR(t) = ΔHbO(t) - ΔHb(t)) across four cortical regions (rows: fNIRS Channel 1 at top through Channel 4 at bottom; columns: Trials 0-3 left to right). (n=18 participants,10 male, 8 female, age= 24-29 years).

Post hoc Tukey comparisons confirmed that T3 differed significantly from all training trials (T0–T2), showing substantially lower values (****p < 0.0001, **p < 0.01). These findings suggest that by the probe phase, the subjects localized their search behavior more precisely around the target.

We measure the differential response (Eq. 8) as the subject is exploring the different regions and generate a spatial map of this differential response (DR) as a function of learning. Fig. 6C shows this differential response normalized to residence time and averaged across all subjects as a function of trials T0–T3 (Each map with its respective scale is shown in SFig. 10). The response maps for training (T0-T2) indicate that the responses surrounding the target region progressively increase relative to the regions outside of where the treasure is located. This is more prominent in long channels’ ch3 and ch4 as compared to ch1 and ch2. While the trajectories determine places that are sampled, the intensities at each of these points are determined by the amplitude of the differential response. The spatial distribution of this amplitude shows a distinct localization in Ch3 and Ch4 and not in Ch1 and Ch2, even though the underlying trajectories are the same and all of them were normalized for the residence time. Thus, indicating that Ch3 and Ch4 activity is correlated with specific retrieval of memory for the location of the treasure. We further quantify this measure by defining contrast measures on absolute differential response maps (see SFig. 8) as provided in the methods.

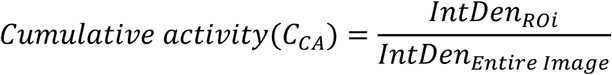

Fig. 7A describes the generation of contrast maps in relation to our device position on the subject’s head. The contrast measured in each of the channels is spatially placed in a geometry corresponding to the device as indicated in Fig. 7B. We find that this contrast progressively increases and fits to a line (Fig. 7C) in Ch3 and Ch4 with a significant slope of 0.37463 ± 0.04356 (t-test for slope > 0 and Adj R = 0.96051) and 0.7399 ± 0.08424 (t-test for slope > 0 and Adj R = 0.96209) for Ch3 and Ch4 respectively. However, this pattern does not hold for Ch1 and Ch2. Their estimated slopes, 0.11047 ± 0.04727 for Ch1 and 0.38913 ± 0.24976 for Ch2 (one-sided t-test for slope > 0), show weaker linear trends, reflected in their lower adjusted R² values (0.59797 and 0.32241, respectively). We also provide such image constructions for other contrasts listed in the methods section of this manuscript in SFig. 9 (B – D).

**Fig. 7.**
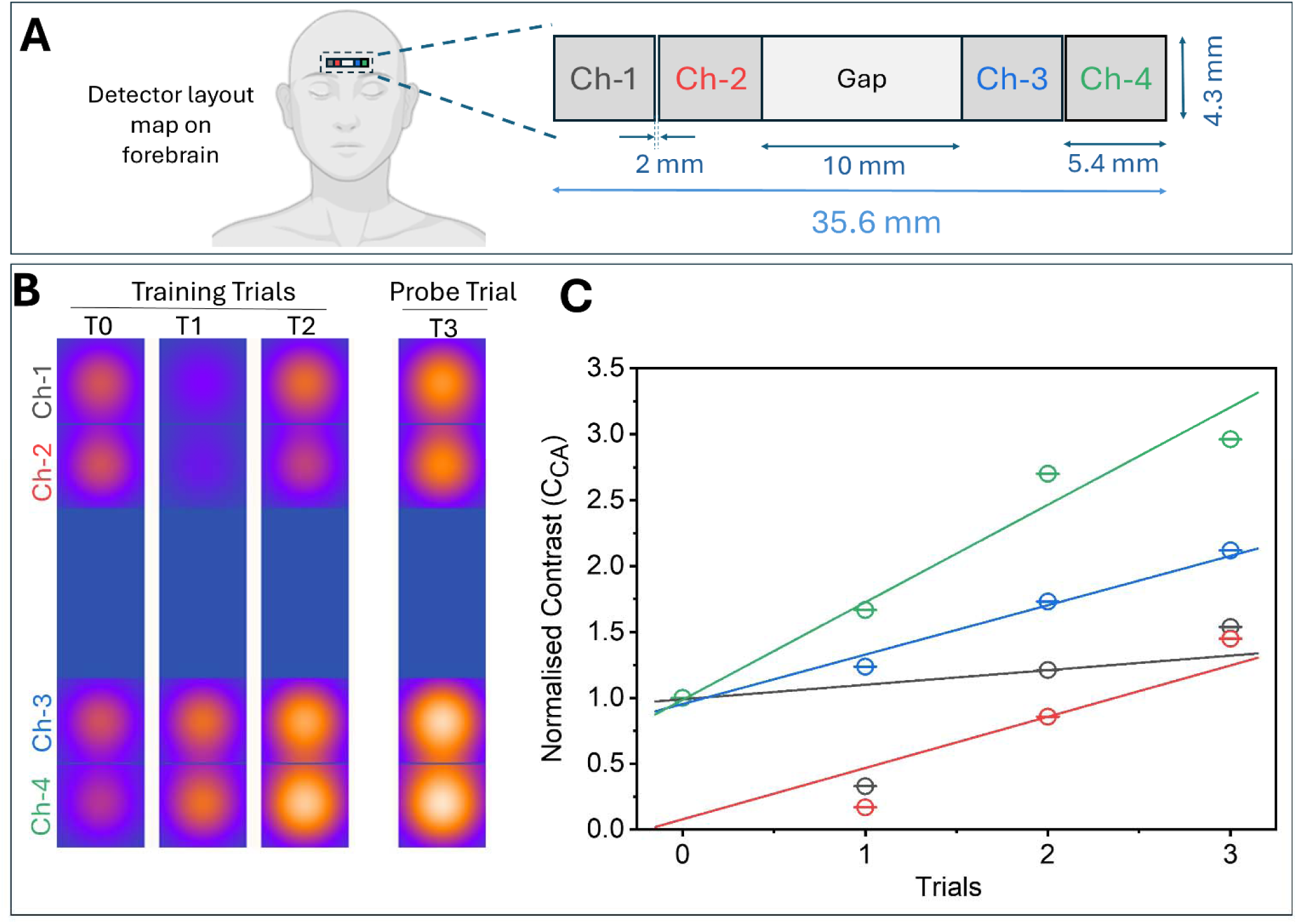
Contrast (C_CA_) maps at subcortical regions. (A) fNIRS channel’s layout on the forehead. *(B)* Contrast maps at different detector positions (Ch1-Ch4) across all trials (T0-T3). ((C) The contrast shows an apparent, progressive increase in Ch3 and Ch4, each following a strong linear trend, with slopes of 0.37463 ± 0.04356 (Adj. R² = 0.96051) and 0.7399 ± 0.08424 (Adj. R² = 0.96209), respectively (one-sided t-test for slope > 0). In contrast, Ch1 and Ch2 exhibit only weak linear increases, as reflected by their smaller slopes (0.11047 ± 0.04727 and 0.38913 ± 0.24976) and lower adjusted R² values (0.59797 and 0.32241).)

## 3. Discussion

Since its initial inception, the NIR region has been used extensively for non-invasive monitoring of hemodynamics. However, the tradeoffs between sensitivity, complexity, versatility and ease of use are limiting factors in using these devices in cognitive assessments. Moreover, the lack of simple procedures to evaluate such devices to estimate the sensitivity further limits the interpretation of the results across the devices. Here we develop a device and characterize its sensitivity in simple, easy-to-implement tests. Our custom-built device was found to be 4.26 times more sensitive (sensitivity calculation is detailed in supplementary information) than a commercial module for detecting oxygenation of hemoglobin. As can be seen from the SFig. 2 C and 2D this increased sensitivity results from lower CV of the signal. We used a two-layer block design that integrates strategically placed detectors and light sources in one block(B1), while another block(B2) has the development board containing a master controller, BLE unit, DAC, and ADC. We reason such a design along minimizes the number of PCB tracks in the device, thereby possibly reducing the noise and overall footprint of our device. We note this is in addition to the inherent differences in the ADCs could have contributed to higher SNR. Use of such a block design further facilitates better coupling of light sources and detectors to the skin/scalp. Thus, it results in an overall improvement in the signal-to-noise ratio. Our two-layer design segregates the optical channel electronics(B1) from data unit(B2), we believe such a design and the use of prefabricated microcontroller unit with a standardized interface in B2 makes it easy to implement and use. This would enable widespread use of the device among the cognitive neuroscientists, clinicians and behavioral scientists for following the oxygen influxes during cognitive tasks.

We functionally validate the increased sensitivity, compactness, and the diverse use of our device. We measure the oxygenation of blood and compare it with a commercial oximeter module and show that not only can we detect smaller changes, but we can also detect them faster, as our sensitivity/time is higher. Owing to this, we could extend the use of our device to measure or monitor the heart rate through a novel and unique method of analysis using Short Time Fourier Transform (STFT).

After the validation of the sensitivity of our device, we used the finger tapping and dynamic contrast task to establish the versatility and functional efficiency of our device in measuring brain activity in motor, sensory and cognitive tasks. In the finger tapping task, we were able to faithfully detect the changes in neuronal activity. The data from a single channel of our device during the finger tapping task showed a clear peak in activity during the tapping period, even in a single subject (averaged over 24 epochs). Averaging across the subjects preserved this characteristic difference between the rest and tapping phase across all the channels (Fig. 3E); however, we see that in Ch3 and Ch4, the decoupling is mixed with an increase in both the amplitude of the concentration change in both Δ[HbO] and ΔHb, yet maintaining the decoupling between the Δ[HbO] and ΔHb. This resulted in an overall increase in the total blood flow into the observation volume. This is in direct contrast to the short channel, where the dominant change is observed in Δ[HbO] amplitude. The Δ[HbO] amplitude sampled by the Ch1 and Ch2 (short channels) increases in the tapping period much more than the ΔHb.

In fact, the ΔHb amplitude of change is more or less constant and is agnostic to tapping and rest phase. In our data the decoupling is observed in all channels, when we normalize the amplitude change of these individual moieties with the change in total concentration of hemoglobin (as can be seen in the lower panels), illustrating the dynamics between Δ[HbO] and Δ[Hb].

A substantial body of experimental work shows that the differential hemoglobin signal, ΔHb_diff_ = Δ[HbO] − Δ[Hb], is a reliable marker of cerebral oxygenation and neural activity. Studies such as ^33,36–39^ demonstrate that the gap between oxygenated and deoxygenated hemoglobin traces the brain’s oxygen exchange, with greater divergence indicating stronger shifts in oxygenation. As a result, simultaneous increases in ΔHbO and Δ[HbT] coupled with a decrease in ΔHb are consistently observed in activated cortical regions.

Increased sensitivity of our device reliably detected the above decoupling with high temporal resolution. In order to measure this decoupling, we construct and use a novel measure, tEIDO, for analyzing the dynamic contrast signals. Since tEIDO is constructed as an instantaneous difference in Δ[HbO] and Δ[Hb], it is akin to measuring the instantaneous change in oxygen influx with high time resolution. We demonstrate that such a measure extracted from our device clearly identifies the activity of the underlying brain region.

The above tasks, which were either motor or sensory tasks, resulted in direct modulation of the corresponding cortices, leading to a detectable fNIRS signal. To demonstrate the utility and versatility of our device, we probed whether a hitherto unknown activation in the brain region could be monitored using a virtual water maze task. Our ability to spatially map and specifically localize activation near the target in long channels demonstrated that this region is involved in memory retrieval. To the best of our knowledge, this is the first direct evidence showing such an activation during memory retrieval. Incidentally, we could also use this information to construct new contrast measures that indicate the differential activation of the four subregions that are possibly involved in this task. Thus, by utilizing BLE as a mode of communication and a single development board for controlling, acquiring, and communicating data, we have been able to simplify the design. Due to its convenient form factor and shape, our device is easy to place on the subject’s head, and we were able to clearly observe the hemodynamics during a diverse set of tasks. Our results showed that the changes deducted from our device have a high SNR and low noise bandwidth. We draw this inference in comparison to the response recorded from the commercial MAX30102 device. Further to enhancing the usability, we show the utility of STFT in measuring the variation in heart rate through time-frequency analysis and graphs. The fact that we could detect changes in activity following single-task activation (for e.g. movement of digits in a hand by a single subject) illustrated the enhanced sensitivity of our device.

We have developed a high-sensitivity BLE-enabled wearable fNIRS device and validated its performance across a battery of standard physiological and neural benchmarks. Our validation tests utilizing conventional tasks, arterial occlusion, and breath-hold in the forearm task, show clear hemodynamics detection. Additionally, we also show that the primary visual cortex activity can be detected in a simple dynamic contrast stimulus presentation. The sensitivity of our device is such that we could easily follow heart rate dynamics using the short-time Fourier Transform. The ability to resolve cardiac pulsations at varying between 0.98 – 1.27 Hz with an accuracy of 0.003 Hz (SFig. 13, Table 1−2). Our ability to resolve and measure heart rate variability with an accuracy of 0.003 Hz demonstrates sensitivity and temporal fidelity. This capability is often limited in bulky, low-sampling commercial fNIRS systems^40–43^. We show that high SNR possessed by our device enables us to follow during a task. To the best of our knowledge, this is the first observation of such task-evoked influxes of oxygen detected from a single subject. Building on these benchmarks, we used the device to follow and study the dynamics of spatial memory in a virtual maze task. On the probe trial, we observed showed intense localized activation in the treasure area. This is prominent in the long channels (ch3 and ch4) that measure the activity from deeper regions.

We reasoned that maze-induced conflict and memory demands would elicit stronger prefrontal oxygenation, consistent with cognitive load theory ^18^. By combining physiological, motor, sensory, and ecological cognitive validations, this study demonstrates that our biosensors not only match benchmark paradigms but also extends wearable fNIRS toward field-ready neuroergonomic applications ^2^.

## 4. Materials and methods

### 4.1 Instrument Design and Fabrication

We designed a wearable four-channel fNIRS system integrating time-multiplexed 740 nm (Marktech Optoelectronics, part No. MTSM0074-843-IR), and 850 nm LEDs Würth Elektronik, part No. 15412085A3060), synchronized silicon PIN photodiodes (VBPW34SR), and a custom transimpedance amplifier (OPA2369AIDGKR, R = 1 MΩ, C = 12 pF, 13.26 kHz bandwidth). All optoelectronics are mounted on a custom designed PCB housed in a silicone-jacketed Velcro patch (Fig. 1B), exposing only source and detector surfaces. We use ESP32-S3-DEVKITC-1 board for LED multiplexing, 10 Hz ADC sampling, and BLE streaming and it is powered by a rechargeable 3.7 V Li-Po battery. The entire device is enclosed in a 3D-printed enclosure that provides programming access and a charging port (Fig. 1C). We also developed the front-layer fNIRS patch in a flexible configuration to enhance skin coupling through conformal adaptation (SFig. 1C). However, data was not collected using this device in the present study.

### 4.2 Data acquisition and transmission

Each LED source was time-multiplexed to eliminate optical crosstalk, remaining active for 27 ms, followed by a 10 ms interval to ensure complete deactivation (SFig. 1A). During each acquisition cycle, a 2 ms delay period preceded the ADC measurement to allow the LED to settle. We found that this delay time is sufficient after measuring the on and off times of the LED using an oscilloscope (data shown in SFig. 2). Following the dual-LED sequence and a 10 ms deactivation interval, a 25 ms baseline period was recorded with all NIR sources switched off. This scheme yields an effective sampling rate of 10 Hz with baseline correction. The timing diagram for this acquisition paradigm is shown in Supplementary Figure 1A. Data acquisition was managed by a Python-based GUI communicating with the fNIRS device over BLE. The device functioned as the BLE server, and the GUI as the client, streaming acquired data to the PC for real-time visualization. The data-collection flowchart and a screenshot of the GUI are shown in. 3(the code is available in our GitHub page).

### 4.3 Materials

a. Optoelectronics and PCB. The complete bill of materials (BOM), circuit schematic, PCB layout, and Gerber files are provided in the project repository.
b. Reference devices for cross-validation. A MAX30102-based pulse-oximeter module (xcluma; MAX30102) and a clinical patient monitor (CONTEC CMS5100; CONTEC Medical Systems) were used to benchmark SpO2 against the developed fNIRS device.
c. Behavioral task software. A custom-made virtual-maze application, developed in the jMonkey Engine, was used to train and test spatial learning and memory. Executable source code is available in the repository.
d. Analysis software. MATLAB R2025a (MathWorks) and OriginPro 2025 (OriginLab).

### 4.4 Participants

A total of 50 healthy volunteers (aged 24-29 years) participated in this study. Each group of participants was assigned to a single experimental paradigm and did not participate in any others. Specifically, 2 male participants were recruited for validation of regional hemodynamics using arterial occlusion and breath-hold tasks, 14 participants (8 male, 6 female) for the motor task experiment, 16 participants (9 male, 7 female) for the visual task experiment, and 18 participants (10 male, 8 female) for the spatial learning and memory task. All experimental protocols were approved by the Institute Human Ethics Committee (IHEC) of the Indian Institute of Science (IISc), Bangalore (approval no.: 04/20.07.2022). Written informed consent was obtained from every participant prior to data collection.

### 4.5 Validation protocols

#### a. Arm occlusion and breath-hold test

We performed simultaneous recordings over 400 seconds using our four-channel fNIRS device, placed on the forearm, and a commercial MAX30102 pulse oximeter module affixed to the index fingertip of the same arm. Vascular occlusion was induced with a CONTEC patient monitor by inflating a blood-pressure cuff on the upper arm; occlusion events were initiated at 90 s and 120 s. At 300 s, subjects executed a 30 s breath-hold. For the MAX30102, we adapted the customized Arduino library for presenting module data to output raw *SpO_2_*% data without internal thresholding; the customized library is available in our repository.

#### b. Motor task: Finger tapping experiment

Participants performed six distinct tapping sequences, each comprising five sub-blocks of 24 taps in total, at a constant 2 Hz pace. Sequences (e.g., 8-6-4-3-3, 3-8-6-3-4, etc.) varied only in the order of sub-block lengths to sustain attention without altering overall cortical load. Each session consisted of 24 cycles of 12-second activation (tapping) followed by 12-second rest (hands flat), for a total run time of 9.6 minutes. A continuous binaural metronome guided movement throughout activation and rest periods, isolating motor-related hemodynamics from auditory-evoked responses. Participants kept their eyes open, minimized head and facial motion, and practiced the task before data collection. At the end of the 24 tapping blocks, a breath-hold protocol (16 s normal breathing, 4 s deep breath, 10 s breath-hold; repeated twice) provided an independent vascular calibration. The task was administered using a custom-written HTML code (provided in the repository). The four-channel biosensor was placed over the area of the left primary motor cortex (all subjects were right-handed) localized by landmarks as described in ^44^.

#### c. Dynamic visual contrast experiment

We designed a block-structured, feature-binding task compatible with fNIRS temporal resolution, preserving the 16s ON/16s OFF timing from ^31^ while incorporating high-contrast, shape-based stimuli ^30^Each trial consisted of a 4 s low-contrast (LC) image, two 4 s high-contrast (HC) images presented in random order, and a final 4 s LC image (total duration = 16 s; images shown in SFig. 12). A session contained 32 trials. A separate baseline (eyes-closed, fixation-cross, and brief breath-hold epochs) was acquired to calibrate cerebral hemodynamics and check the detector’s coupling to the scalp. This task was also administered using a custom-written HTML code (provided in the repository). The four-channel fNIRS biosensor was placed over the occipital cortex (BA 17-19) using 10-20 EEG landmarks. Optodes were housed in Velcro headbands for stable skin coupling.

### 4.6 Cognitive application: Spatial navigation task in a virtual environment

We implemented a custom on-screen spatial navigation task modeled after the Morris water maze to assess participants’ ability to associate visual landmarks with a hidden treasure. The maze is divided into four imaginary quadrants (Q1-Q4). The hidden treasure was located in Q2, as shown in Fig. 6A. In our laboratory-developed virtual environment, each subject explored a maze marked by three distinct cues and was allotted 60 seconds per trial to locate the concealed treasure. During the first three acquisition trials, failure to find the treasure within 60 s triggered a flag at its position.Subjects were instructed to observe and memorize the treasure’s location relative to the visual cues. Subjects need to hit the flag post to end the current trial and go to the next trial. The treasure was removed in the subsequent probe trial (trial 4), and participants again searched for 60 s without the knowledge of the treasure being removed. Successful learning was defined by a statistically significant increase in dwell time within the former treasure zone, indicating recall of its location through cue association. Simultaneously, we recorded prefrontal cortical hemodynamics using our wearable four-channel fNIRS device.

### 4.7 Data analysis and cortical mapping

#### 4.7.1 Calculating the concentration change of Molecules

The dataset from the device consists of photodiode currents that have been converted into voltage through amplification for different wavelengths of light, recorded over a specific period. This measured voltage is proportional to a particular wavelength’s light intensity (I) falling on a photodiode-sensitive area. Keeping the source intensity or incident intensity (*I*_0_) constant over time, we calculate the change in optical density (Δ*OD*) by taking two detected intensity values *I*_1_ and *I*_2_ at two time points *t*_1_ and *t*_2_ respectively for a particular wavelength(λ). We can derive Δ*OD* from MBLL as follows

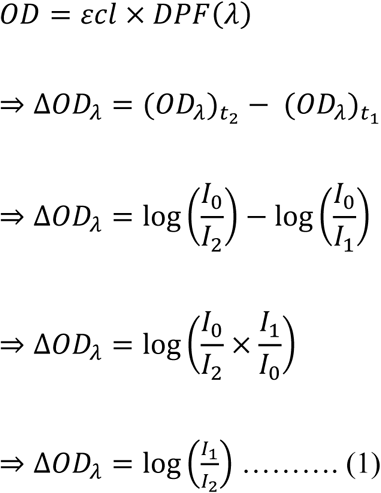

where,

*I_o_ = Incident Intensity*

*I_1_ = Detected Intensity at t_1_*

*I_2_ = Detected Intensity at t_2_*

*ε = Extinction Coefficient;*

*l = source – Detector distance*

Thus, for two wavelengths, MBLL is given as

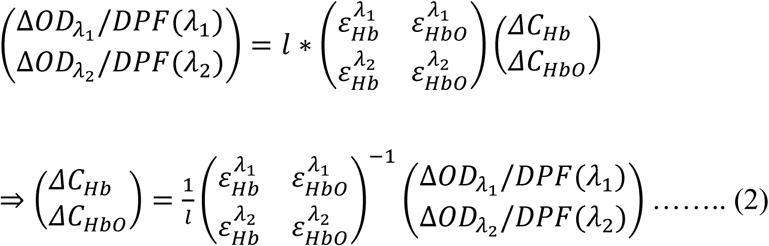

Where,

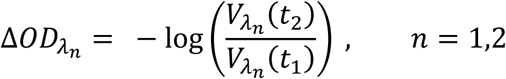

Δ*C*_*HbO*_ = change in concentration of Oxy-Hemoglobin

Δ*C*_Hb_ = change in concentration of Deoxy-Hemoglobin

Therefore, using this approach, the change in the concentration can be calculated from the dataset containing raw voltages. We adopted *DPF(740nm) = 6.29, DPF(850nm) = 5.6(extrapolated)* from ^45^ and 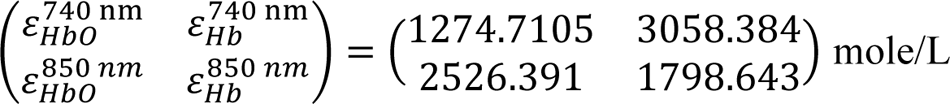 from^46^.

#### 4.7.2 Residence time heatmap

We used each participant’s recorded Cartesian (X, Y) trajectories within the maze to generate a residence-time heatmap across all subjects using a vector analysis software (Meenakshi et al., 2022. Each resulting heatmap was peak-normalized to standardize its dynamic range. Moreover, we derived two region-of-interest (ROI) masks from the normalized heatmaps: (1) a “maze-wide” mask by thresholding all pixels with values above zero, defining the sampled space, and (2) a “treasure-specific” mask by applying the same threshold only within the known treasure location for further analysis.

#### 4.7.3 Activity and differential response heatmap

From the combined spatial (XY coordinates)-activity data (Δ[HbO] and Δ[Hb]), we generated separate activity heatmaps for Δ[HbO] and Δ[Hb] across subjects. Each activity heatmap was then normalized by the previously computed residence-time heatmap (before peak normalization of the residence heat maps) to correct for sampling density and then applied a spatial Gaussian filter with σ=50. We split the normalized activity heatmaps into positive (pixel values > 0) and negative (pixel values < 0) halves, followed by peak normalization to standardize their dynamic ranges, as shown in SFig. 5.

We also generated differential response maps *(DR(t) = ΔHbO(t) − ΔHb(t))* by subtracting (before residence time normalization) the activity map of Δ[Hb] from Δ[HbO] and normalizing by the residence time heatmap and applying a Gaussian filter with σ=50.

#### 4.7.4 Contrast measurement metrics

We have quantified the change in heatmap contrasts towards the treasure location (ROI) relative to the entire maze (Entire Image). We have defined four metrics (Eq. 3-6) to capture the contrast.

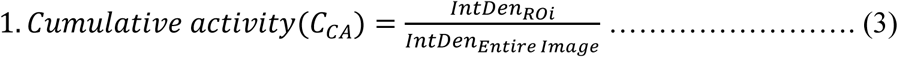

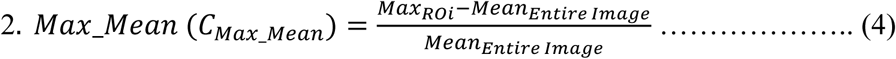

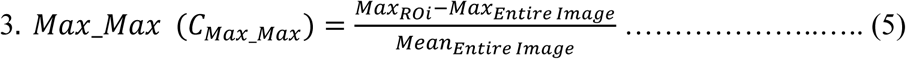

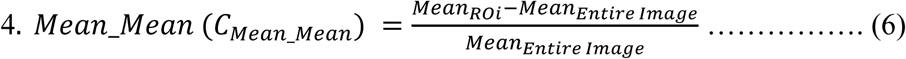

For the activity heat maps of Δ[HbO] and Δ[Hb] (both positive and negative, as mentioned in section 4.7.3), we used the ROI masks defined in section 4.7.2 and computed the metrics within each trial to define “goal-specific” (around the hidden treasure location) neural activity change in different brain regions covered by each detector (SFig. 6-7).

To compute the above metrics to capture the contrasts in differential response maps, we used the absolute maps. Finally, these quantitative metrics were mapped to the device’s detector layout to assign each measure to its corresponding detector location, and contrast maps were produced as described in Section 4.7.5.

#### 4.7.5 Subcortical Mapping

We modeled each fNIRS detector as a 60×60-pixel tile (corresponding to a 5.4 mm × 4.3 mm physical sensor area). Detectors D1-D4 were arranged in a planar array with a 10 mm gap between the active edges of the adjacent detector pairs (D1-D2, 10 mm-D3-D4), accurately reproducing the device’s geometry and physical dimensions. Within each 60×60 detector tile, we defined the central 10×10-pixel region to assign the calculated measures described in Section 4.7.3 as the raw intensity. To eliminate negative intensity values and stabilize downstream processing, all pixel values were offset by adding twice the absolute value of the global minimum across that detector’s measurement window:

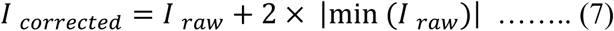

A two-dimensional Gaussian filter (σ = 20 *pixels*) was applied independently to each detector’s 60×60 tile, producing a smoothly varying intensity map. Finally, a standardized lookup table (LUT) was overlaid to smooth detector signals for mapping goal-specific hemodynamic activity.

## Funding Support

The work was supported by funding received from ICMR with grant ID: IIRP – 2023-4723 CPDA to BJ and IISc Intramural funds.

## Notes

### Competing Interest Statement

The authors have declared no competing interest.

