## Supplementary Figures and Tables for "Tracking instances of task evoked oxygen influxes using a novel sensitive BLE enabled wearable fNIRS device identifies mPFC role in spatial memory"

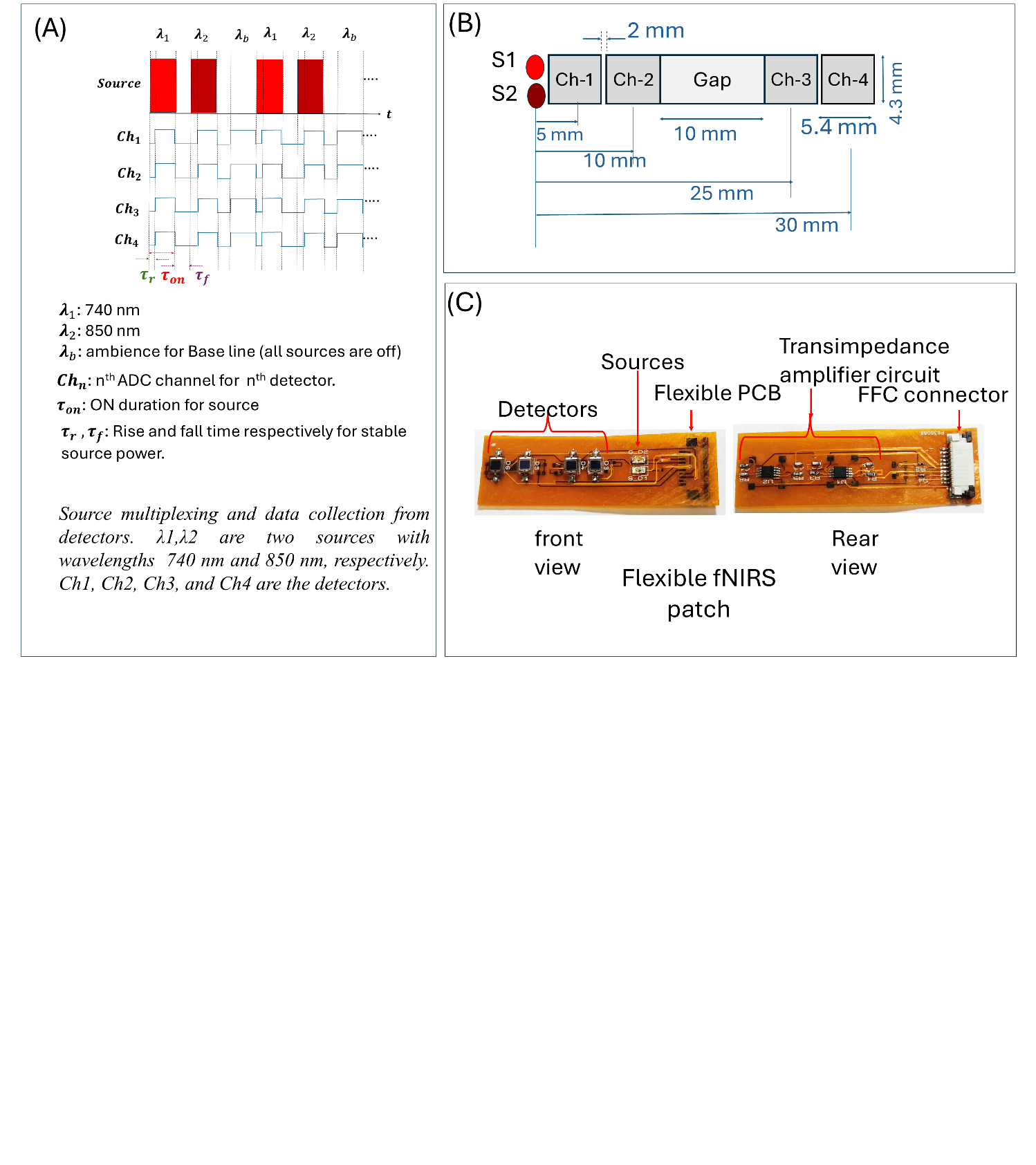

Fig. 1. (A) The data acquisition paradigm shows the time multiplexing of sources and reading through ADC pins. (B) The fNIRS patch's physical layout shows the location configuration of the source and detectors at certain distances. (C) The fNIRS patch is built with flexible construction (flexible PCB).

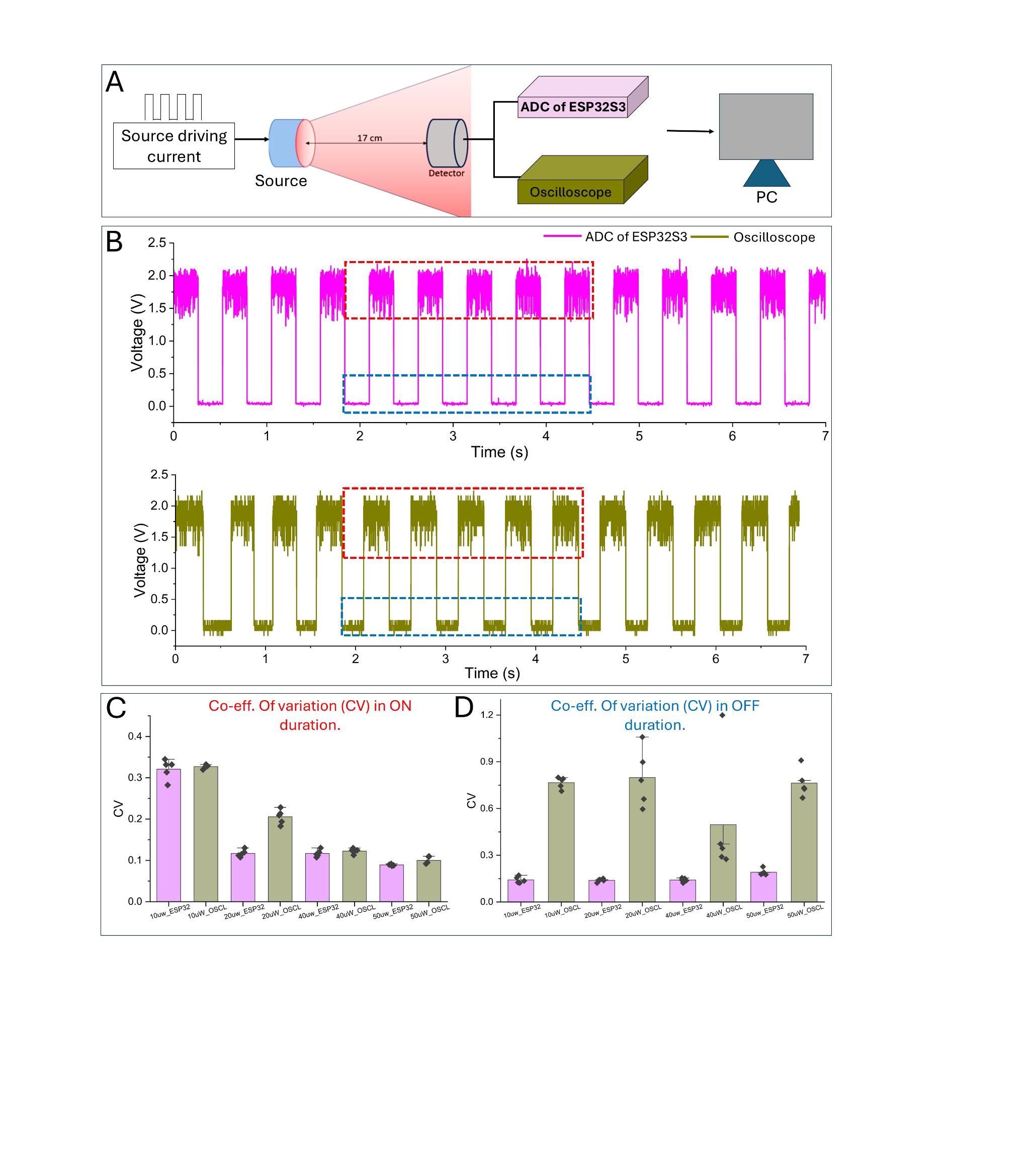

Fig. 2. Simultaneous acquisition of detector responses using the ESP32-S3 and an oscilloscope (SIGLENT SDS1104X-E). (A) Output voltages from the detector were recorded concurrently by the ESP32-S3 and the oscilloscope. (B) Detector output traces captured by the ESP32-S3 (top) and the oscilloscope (bottom) under 50 µW incident light over a 7-second period. (C) Coefficient of variation for five ON-state voltage segments (highlighted with the red dashed box in panel B) measured by the ESP32-S3 and oscilloscope for different incident optical power. (D) Coefficient of variation for five OFF-state (baseline) voltage segments (highlighted with the blue dashed box in panel B) measured by the ESP32-S3 and oscilloscope for different incident optical power.

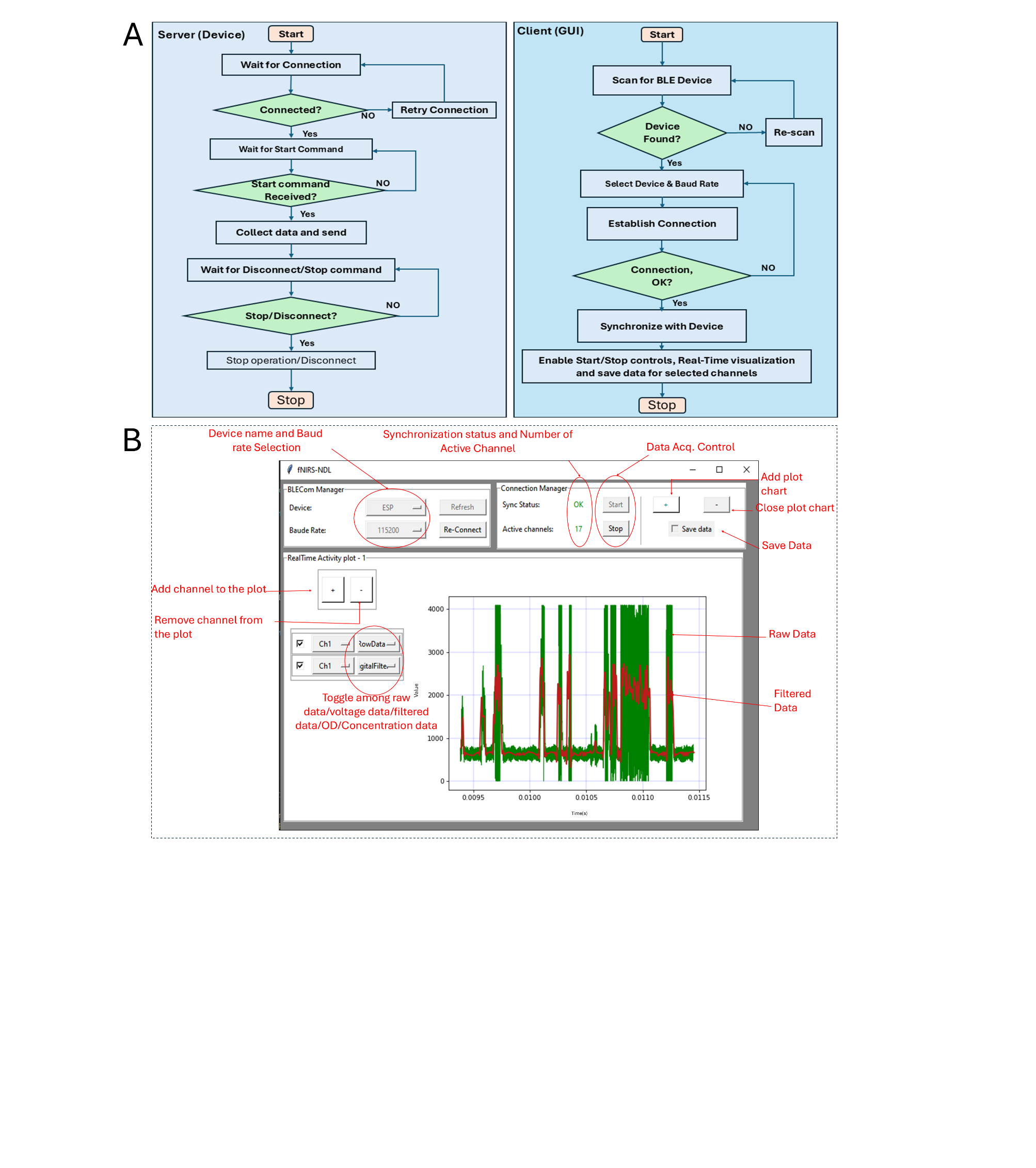

Fig. 3. (A) Data-collection flowchart for the server (the fNIRS device) and the client (GUI running on PC). (B) A screenshot of the graphical user interface (GUI showing different control options.

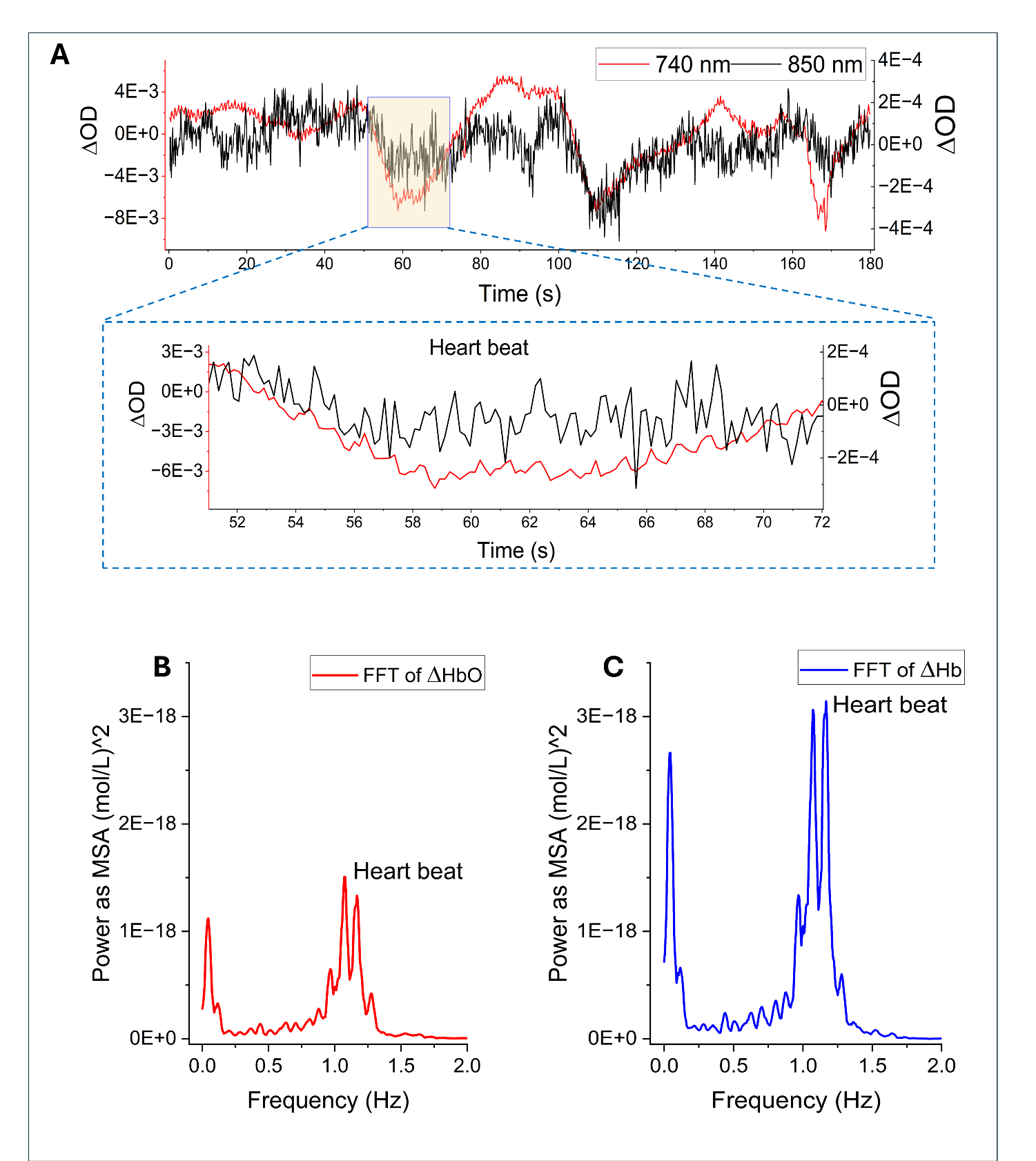

Fig. 4. (A) Change in Optical density of 850 nm (black ) and 740 nm (red) during breath-hold test. (B) and (C) Power spectrum of calculated ∆[HbO] (red) and ∆[Hb] (blue) respectively.

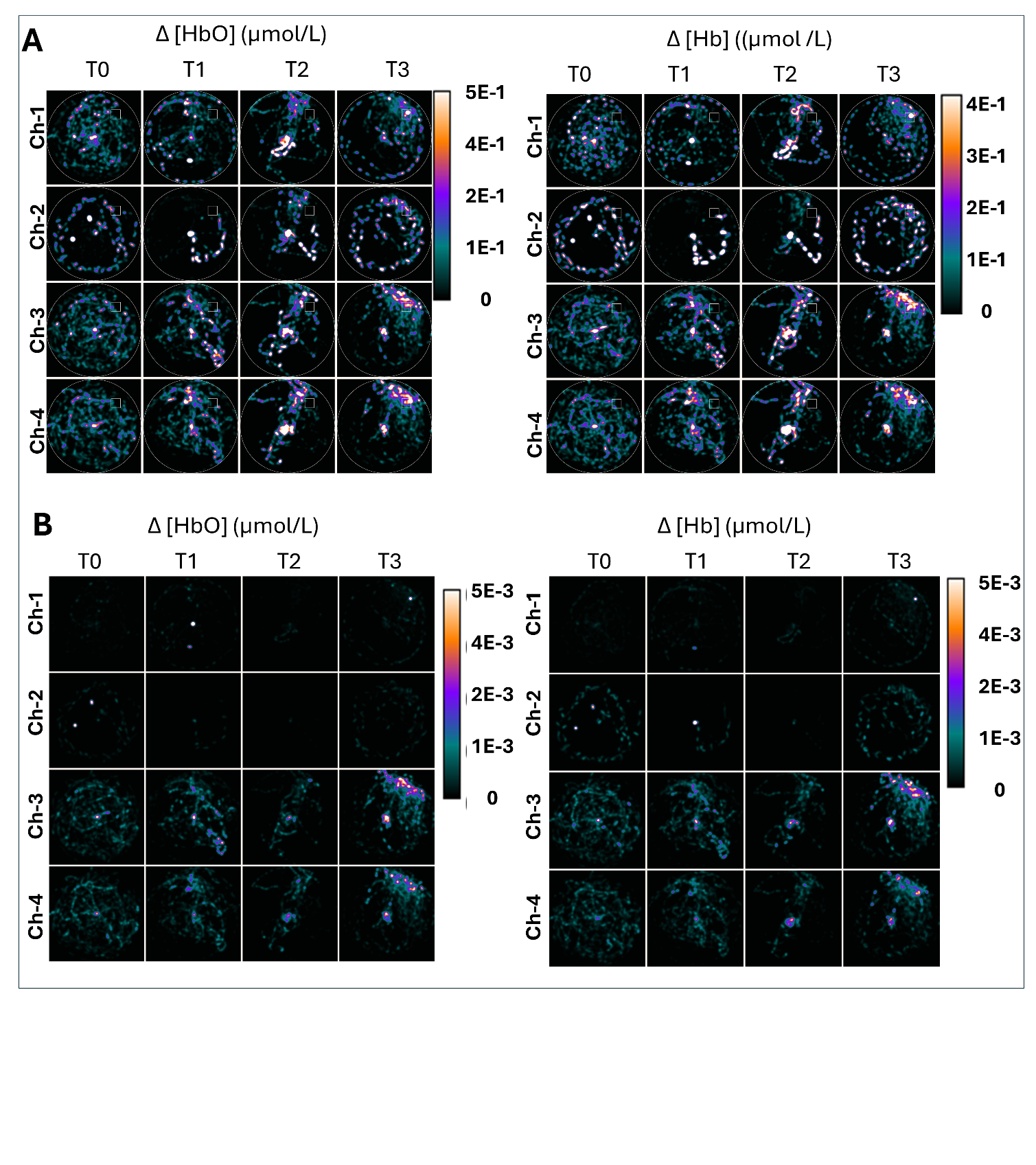

*Fig. 5. Residence time normalized hemodynamic responses across four fNIRS channels (rows: Channel 1 at top through Channel 4 at bottom; columns: Trials 0-3 left to right) across all subjects (n=18). The heatmap displays spatial-temporal activity in terms of Δ[HbO] (oxyhemoglobin) and Δ[Hb] (deoxyhemoglobin). (A) Activity heat map with positive half (Δ[HbO] and Δ[Hb] > 0). (B) Activity heat map with negative half (Δ[HbO] and Δ[Hb] < 0). Inverted LUT is used for (B).*

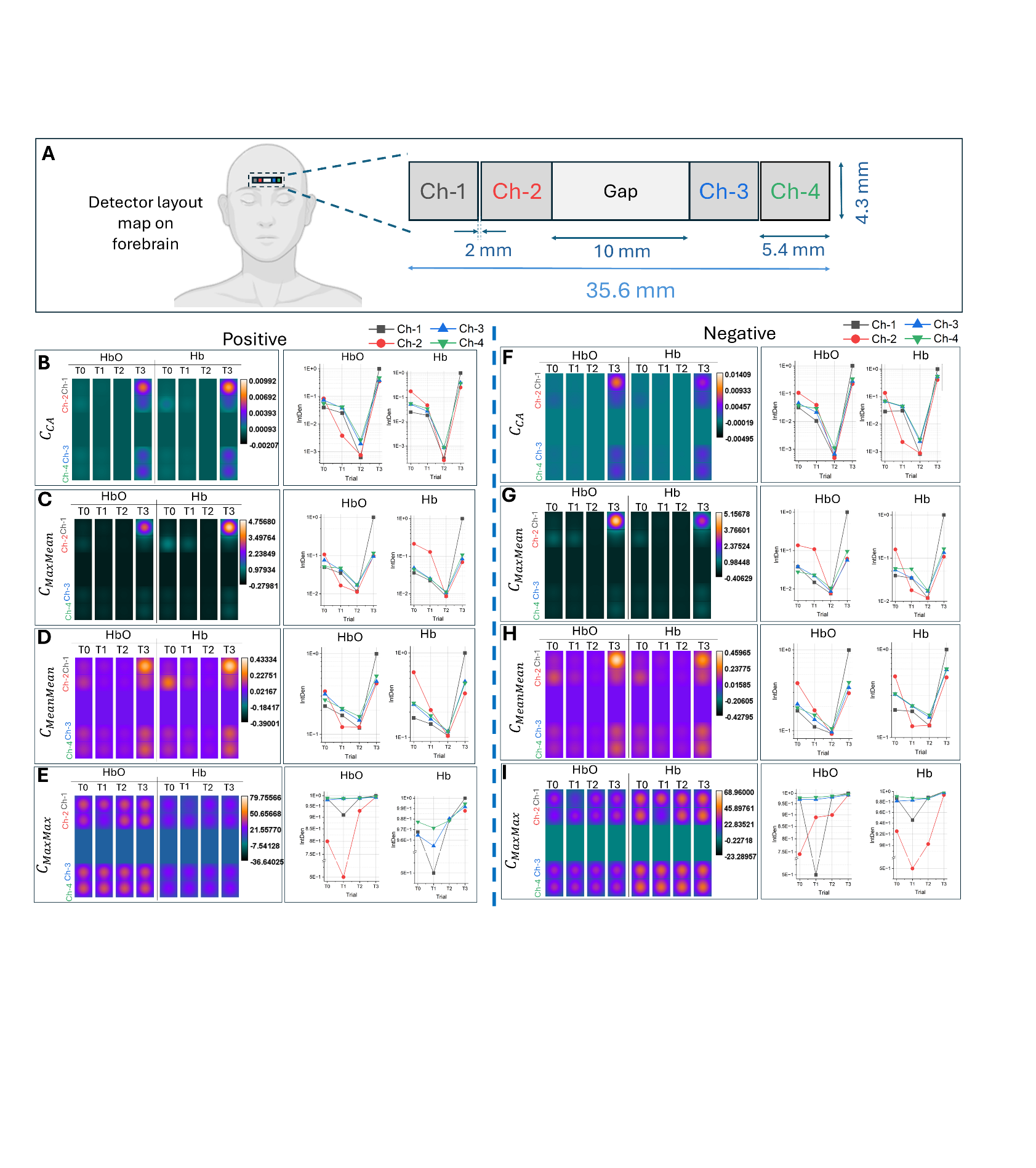

*Fig. 6. Goal-specific hemodynamic activity mapping across trials. (A) fNIRS channel’s layout on the forehead. (B) Trial-wise ∆[HbO] and ∆[Hb] responses at different detector positions on the forebrain, computed as the ratio of integrated activity within the platform region to the total sampled area (left). Corresponding integrated densities from each detector are shown as color-coded line plots: Ch-1 (black), Ch-2 (red), Ch-3 (blue), and Ch-4 (green) (right). (C) Trial-wise ∆[HbO] and ∆[Hb] responses computed as the difference between the maximum value within the platform region and the global maximum across the sampled area, normalized by the global mean (left). Corresponding integrated densities for each detector are plotted on the right using the same color coding. (D) Hemodynamic activity computed as the difference between the platform maximum and the global mean, normalized by the global mean (left). Detector-wise integrated density plots across trials are shown on the right with consistent color coding. (E) Hemodynamic activity computed as the difference between the platform mean and the global mean, normalized by the global mean (left). The right panel shows integrated density within each channel across trials, using the same color scheme.*

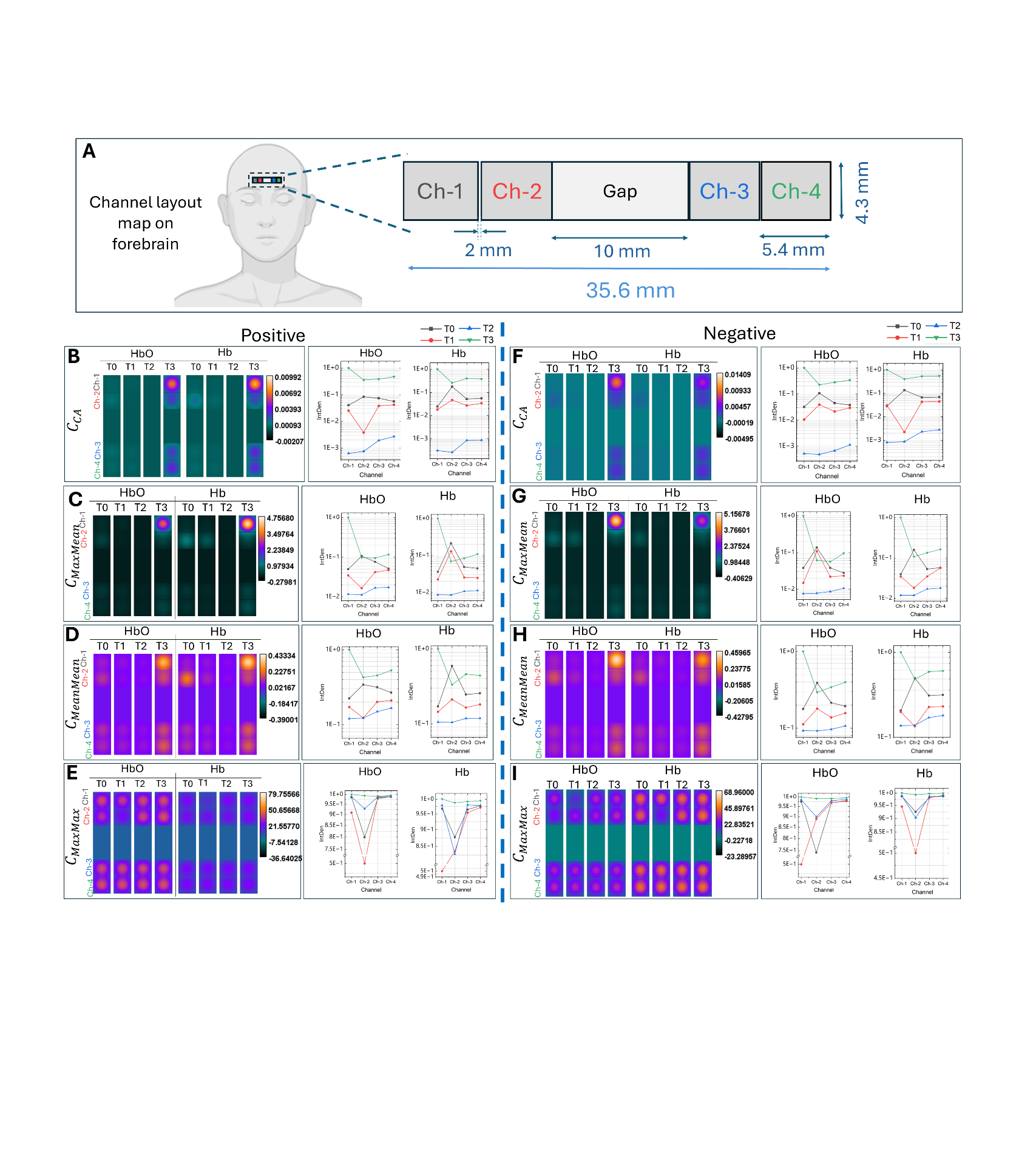
Fig. 7. Goal-specific hemodynamic activity mapping across channels. (A) fNIRS channel’s layout on the forehead. (B) Channel-wise ∆[HbO] and ∆[Hb] responses in different trials, computed as the ratio of integrated activity within the platform region to the total sampled area (left). Corresponding integrated densities from each detector are shown as color-coded line plots: T0-black (trial 0), T1-red (trial 1), T2-blue (trial 2), and T3-green (trial 3) (right). (C) Channel-wise ∆[HbO] and ∆[Hb] responses computed as the difference between the maximum value within the platform region and the global maximum across the sampled area, normalized by the global mean (left). Corresponding integrated densities for each detector are plotted on the right using the same color coding. (D) Hemodynamic activity computed as the difference between the platform maximum and the global mean, normalized by the global mean (left). Channel-wise integrated density plots across trials are shown on the right with consistent color coding. (E) Hemodynamic activity is computed as the difference between the platform and global mean, normalized by the global mean (left). The right panel shows integrated density within each channel across trials, using the same color scheme.

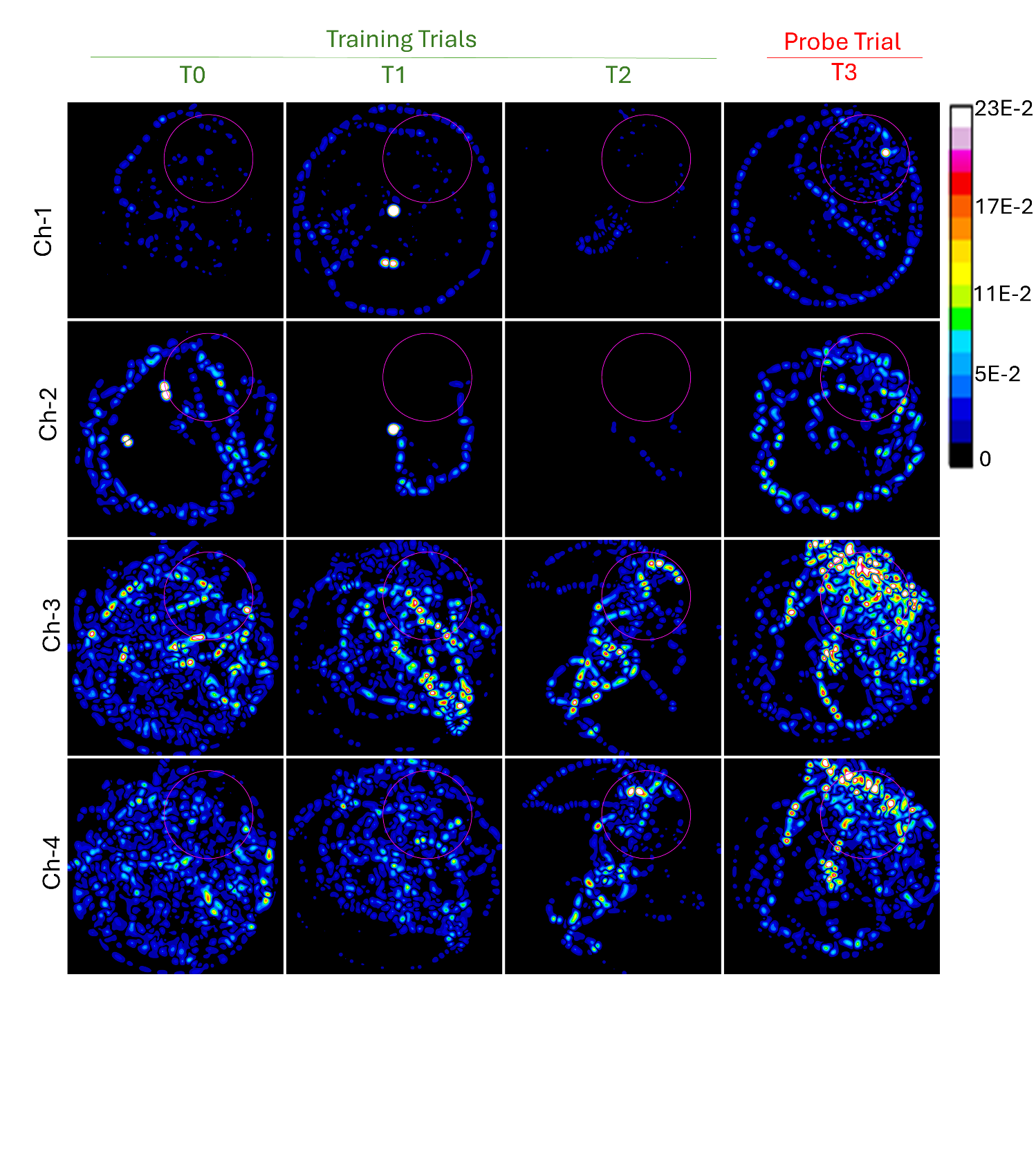

Fig. 8. Absolute differential response (│$DR\left( t \right)=\Delta[HbO\left( t \right)]- \Delta[Hb\left( t \right)]│$) during the task. Rows correspond to Channels 1–4 (ch-1 to ch-4), and columns represent the four trials, where T0–T2 denote the training trials and T3 is the probe/testing trial. The magenta circle represents a zone around the hidden treasure.

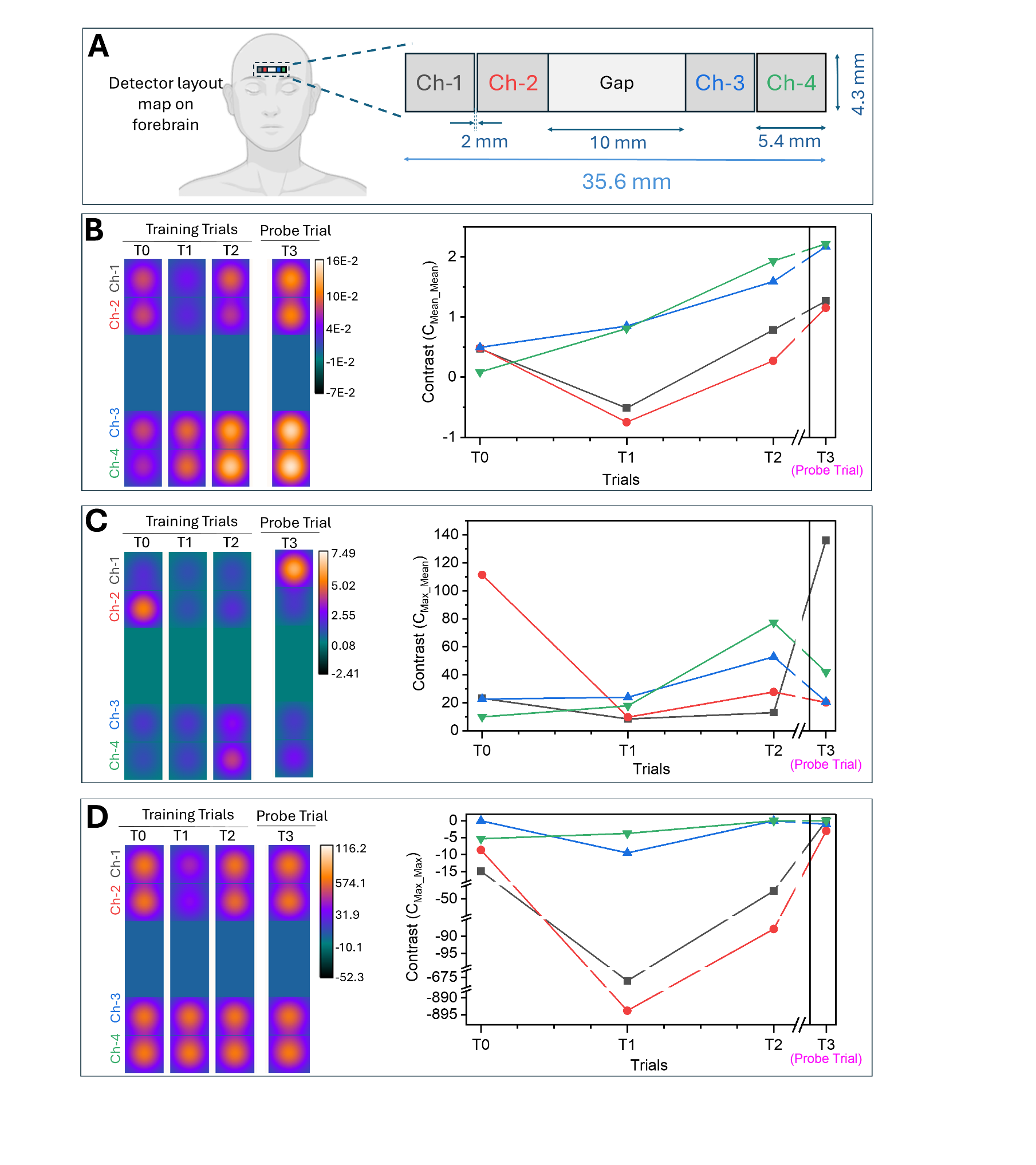

Fig. 9 Contrast maps at subcortical regions. (A) fNIRS channel’s layout on the forehead. (A) Contrast maps computed with the C_Mean_Mean_ metric. (B) Contrast maps computed with the C_Max_Mean_ metric. (C) Contrast maps computed with the C_Max_Max_ metric.

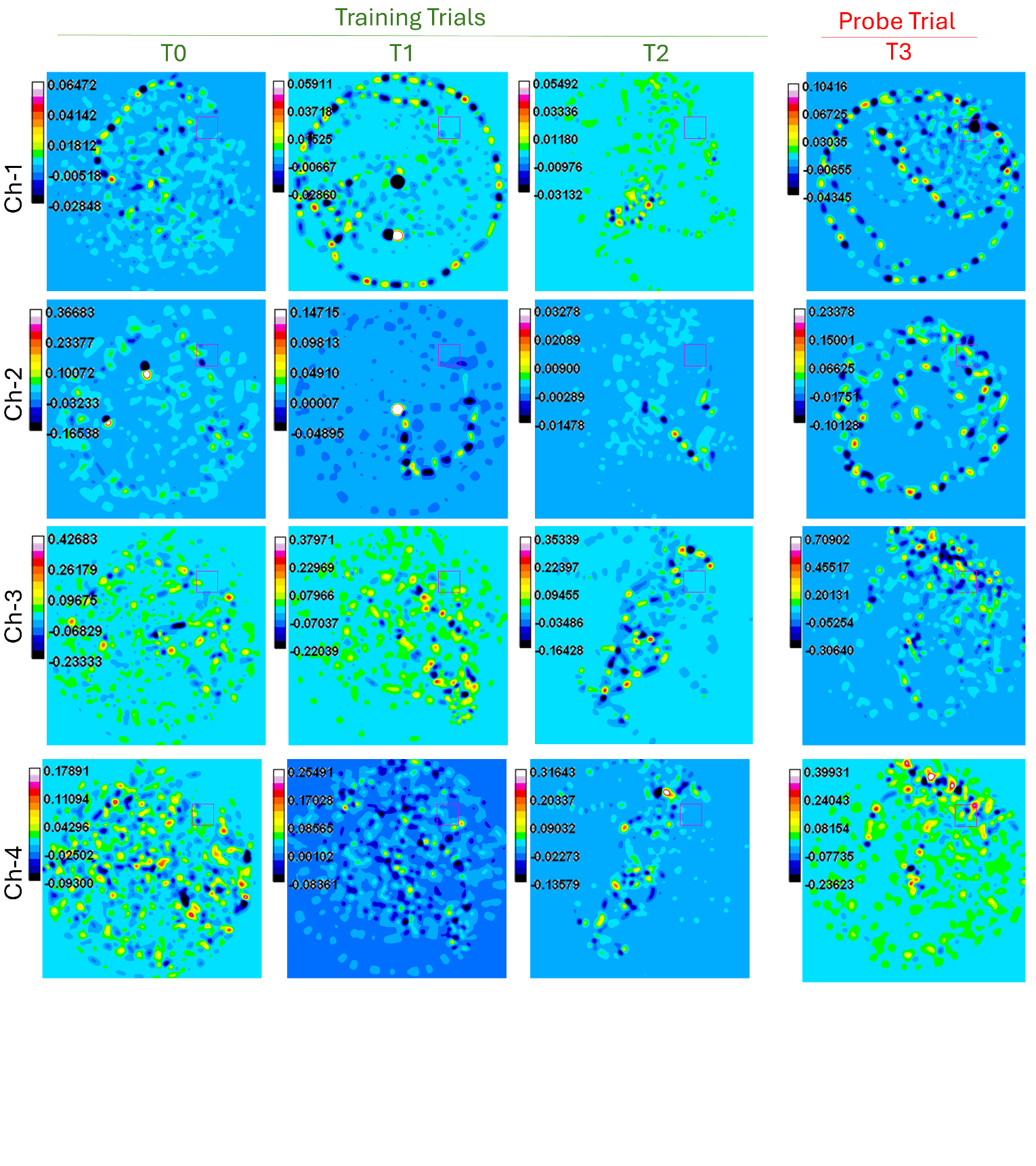

Fig. 10. Differential response ($DR\left( t \right)=\Delta[HbO\left( t \right)]- \Delta[Hb\left( t \right)]$)with an individual contrast map. Rows correspond to Channels 1–4 (ch-1 to ch-4), and columns represent the four trials, where T0–T2 denote the training trials and T3 is the probe/testing trial. The magenta box represents the hidden treasure location.

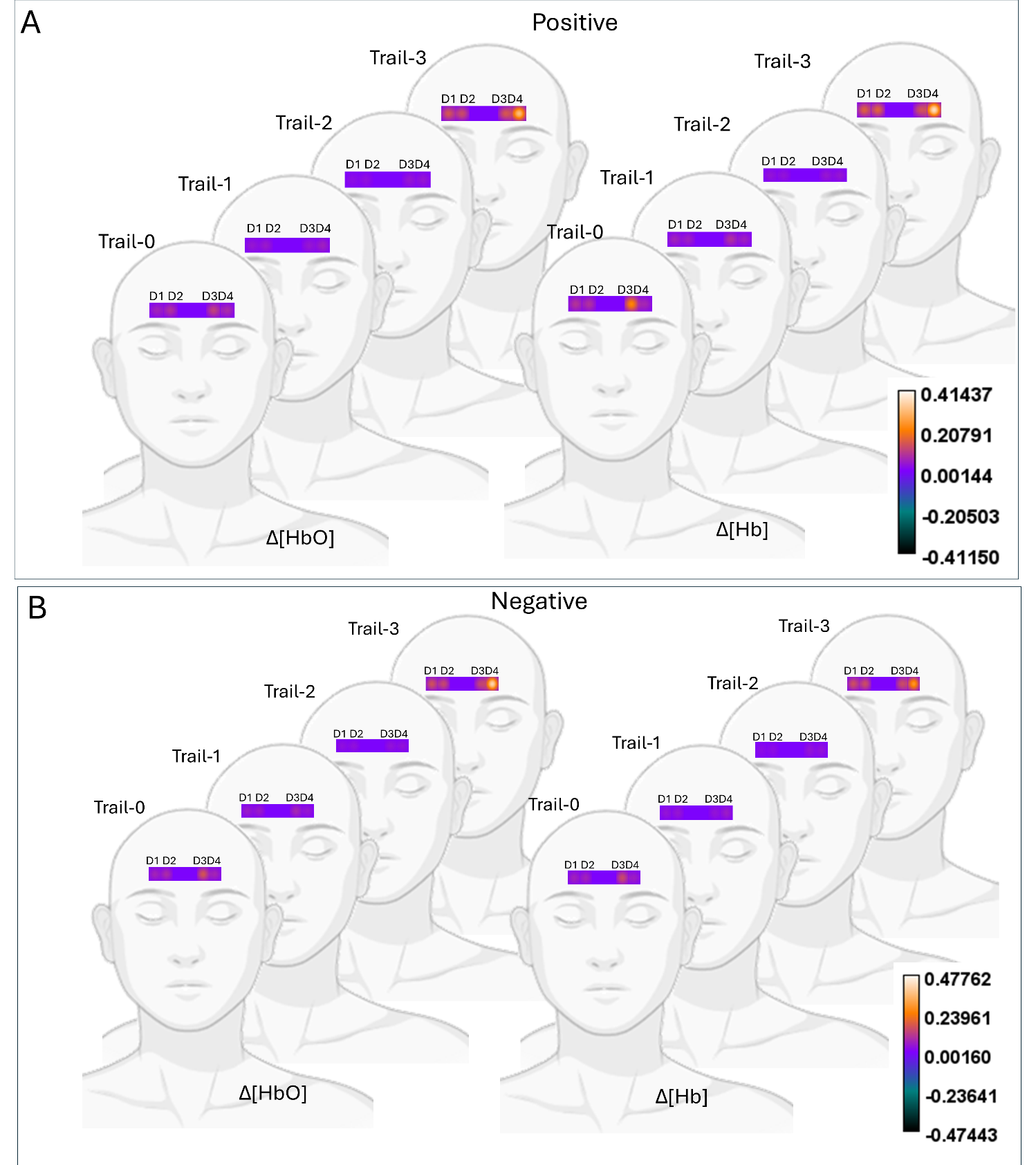

Fig. 11. Goal-specific hemodynamic mapping of the forebrain. (A) and (B) shows goal-specific activity ($C_{Mean\_Mean})$ Mapping for HbO and Hb, respectively.

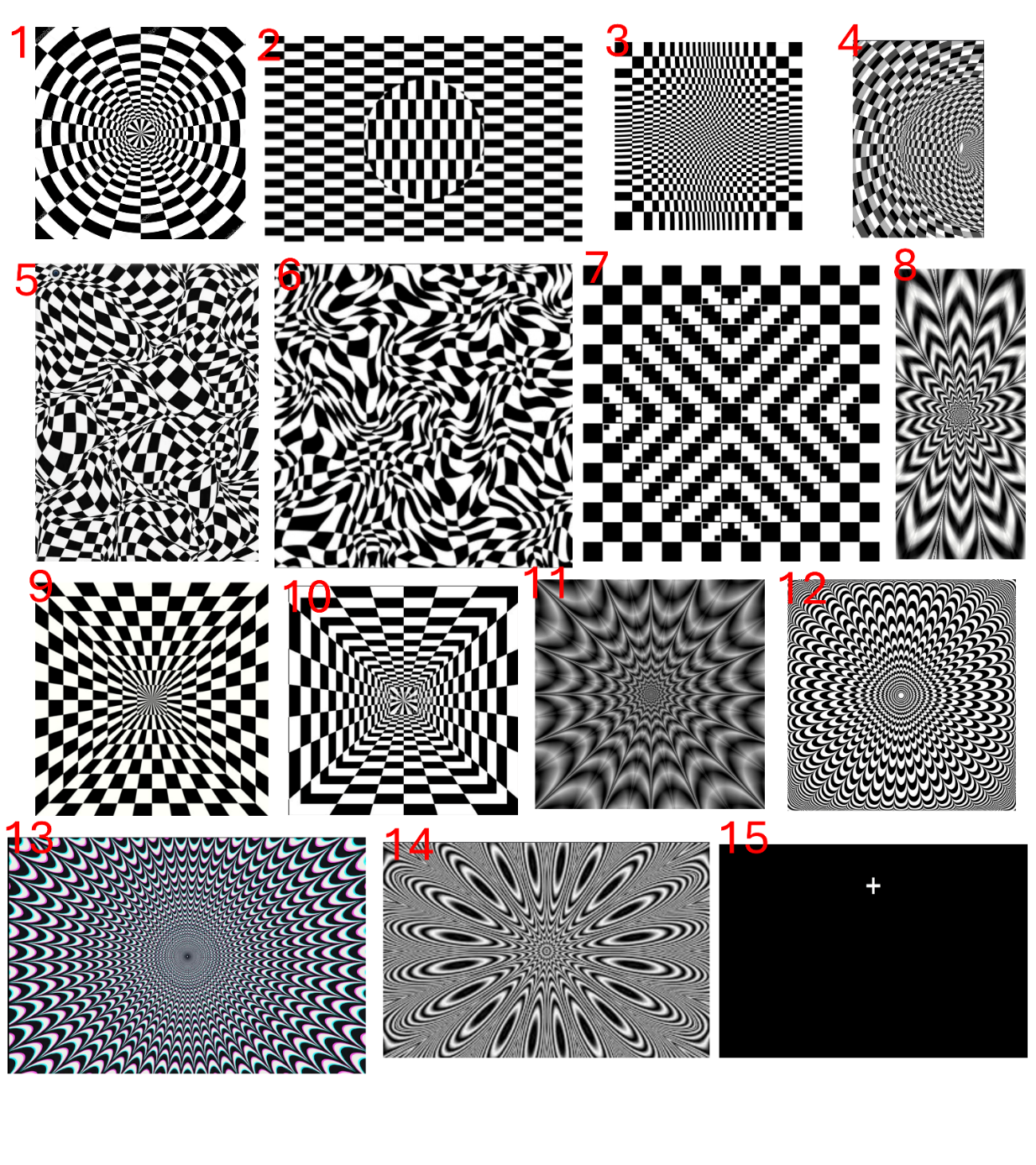

Fig. 12. Images presented as high contrast stimulus (1-14) and low contrast (15).

**Calculation of the Sensitivity:**

$$S=\frac{V_{p}}{\sigma_{b}\times\tau_{b}}$$

$V_{p} =Normalized mean peak value (for our fNIRS device=4.18744 and for Oximeter module=2.63484).$

$$\sigma_{b}=standard deviation of the baseline (for our fNIRS device=0.07248, Oximeter module= 0.19433)$$

$$\tau_{b}= baseline duration (61.13537 seconds (294.2261 seconds to 233.09073. seconds))$$

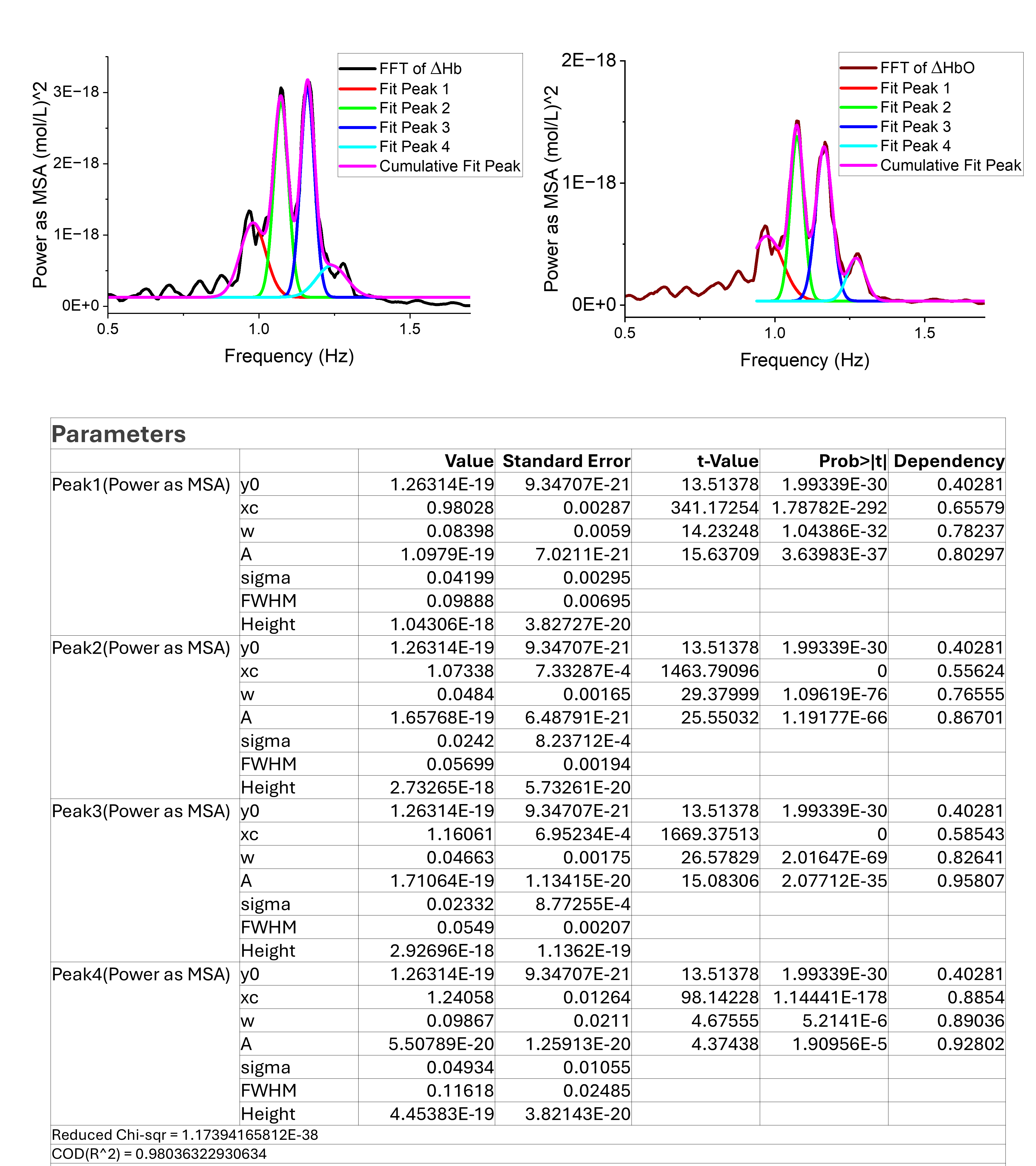

Fig. 13. Curve fits applied to the dominant peaks of the Δ[Hb] (left) and Δ[HbO] (right) FFT spectra.

**Table-1:** Estimated parameters from the curve-fitted FFT spectrum of Δ[Hb]

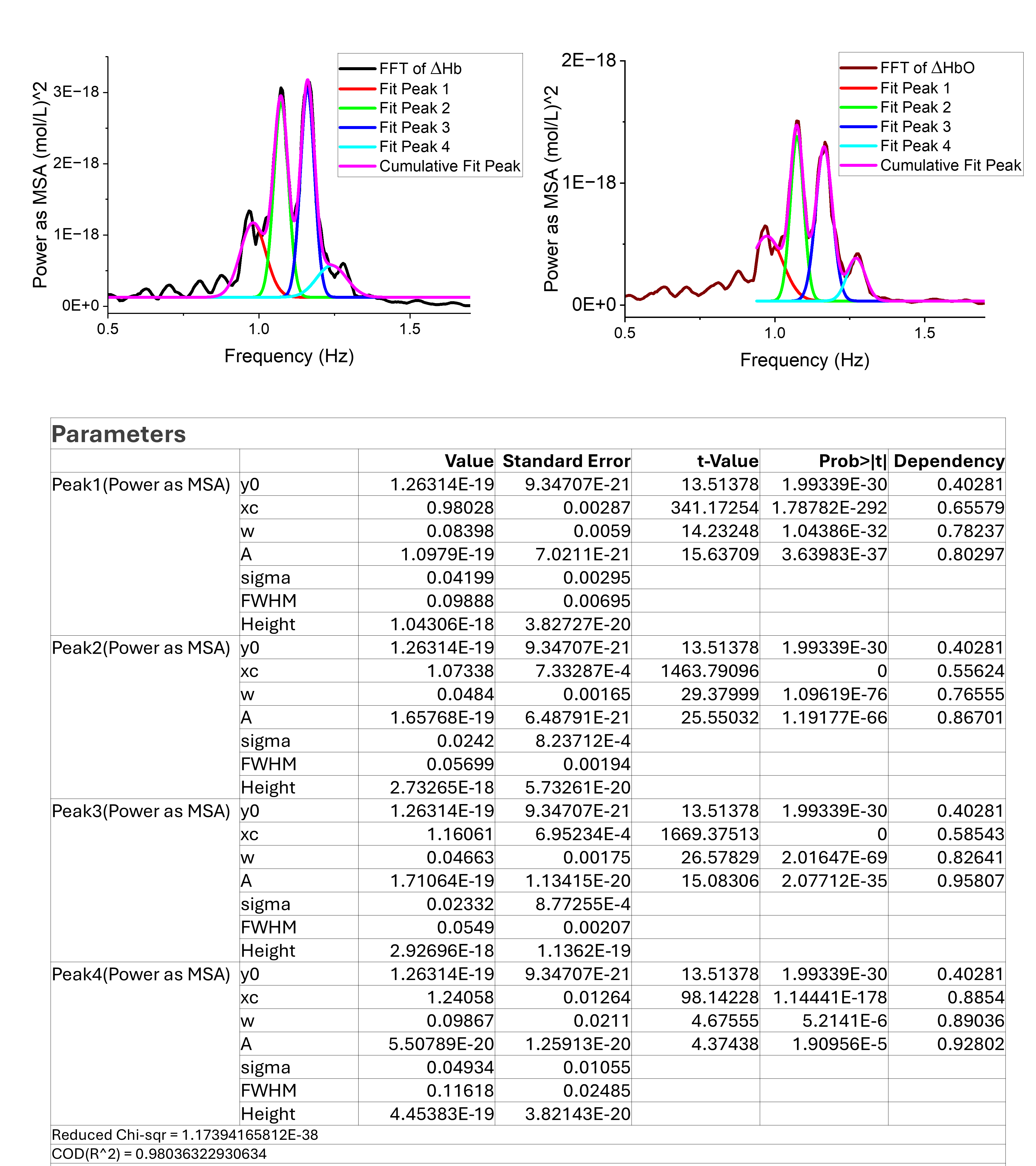

**Table-2:** Estimated parameters from the curve-fitted FFT spectrum of Δ[HbO]

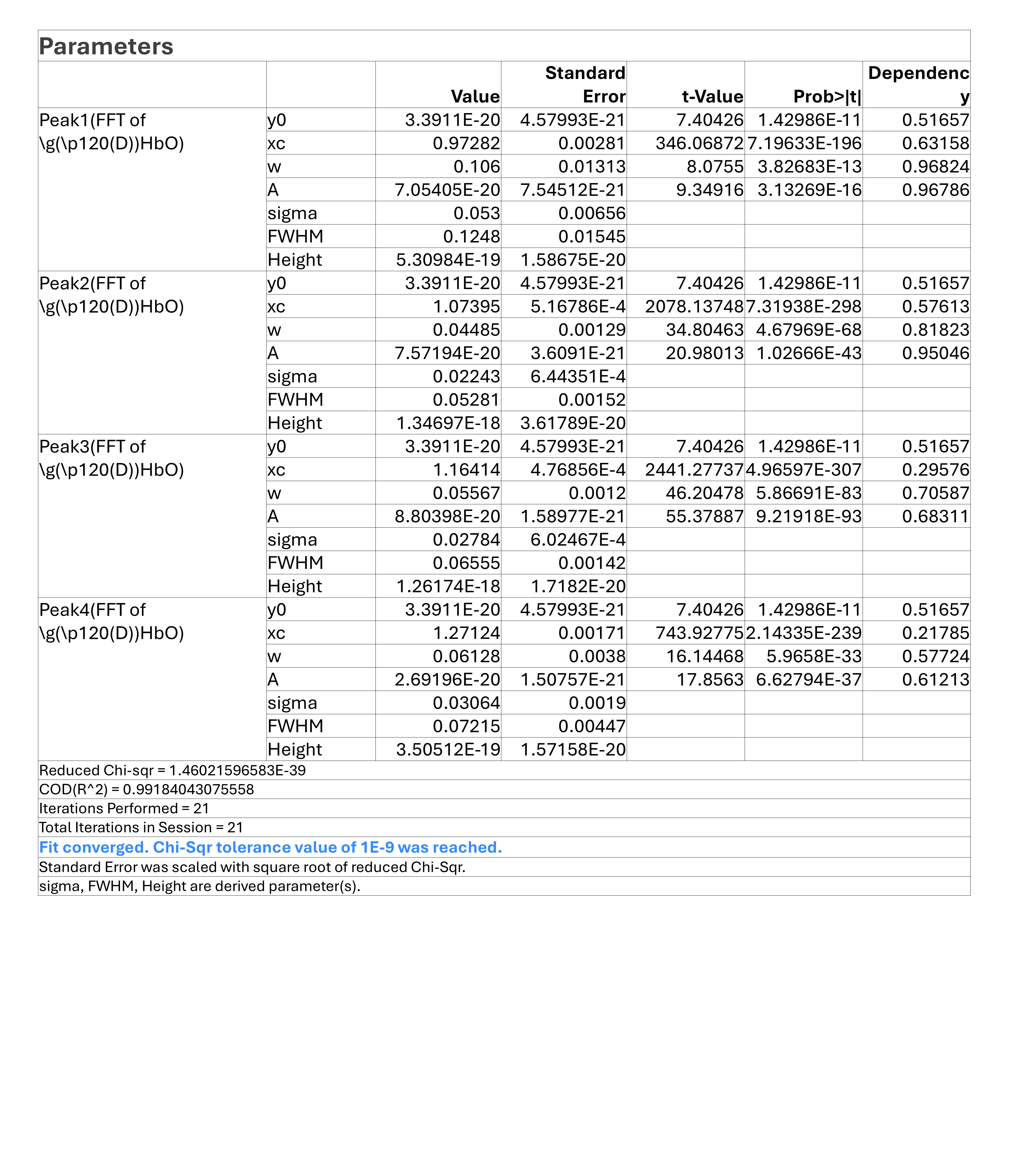

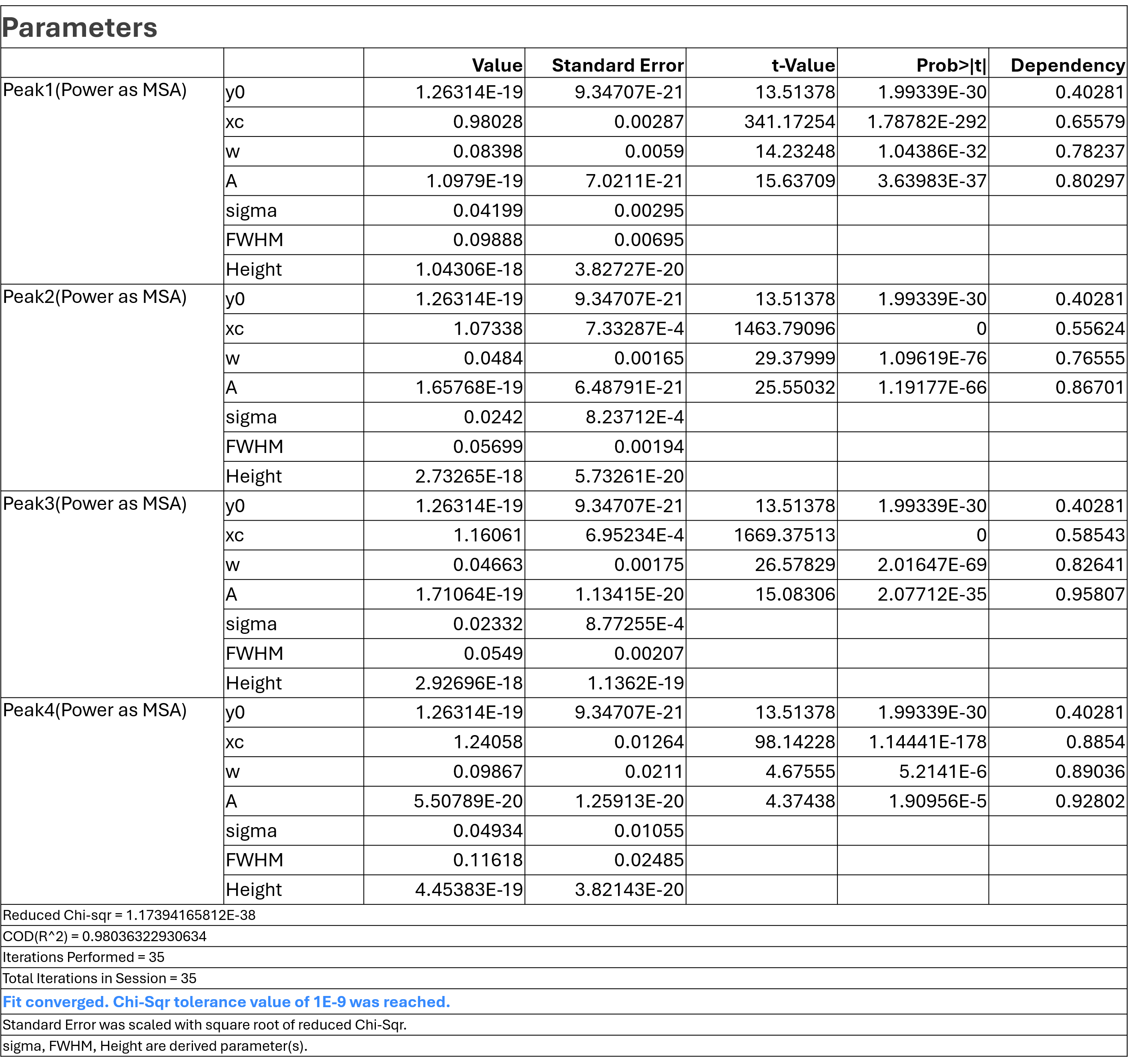

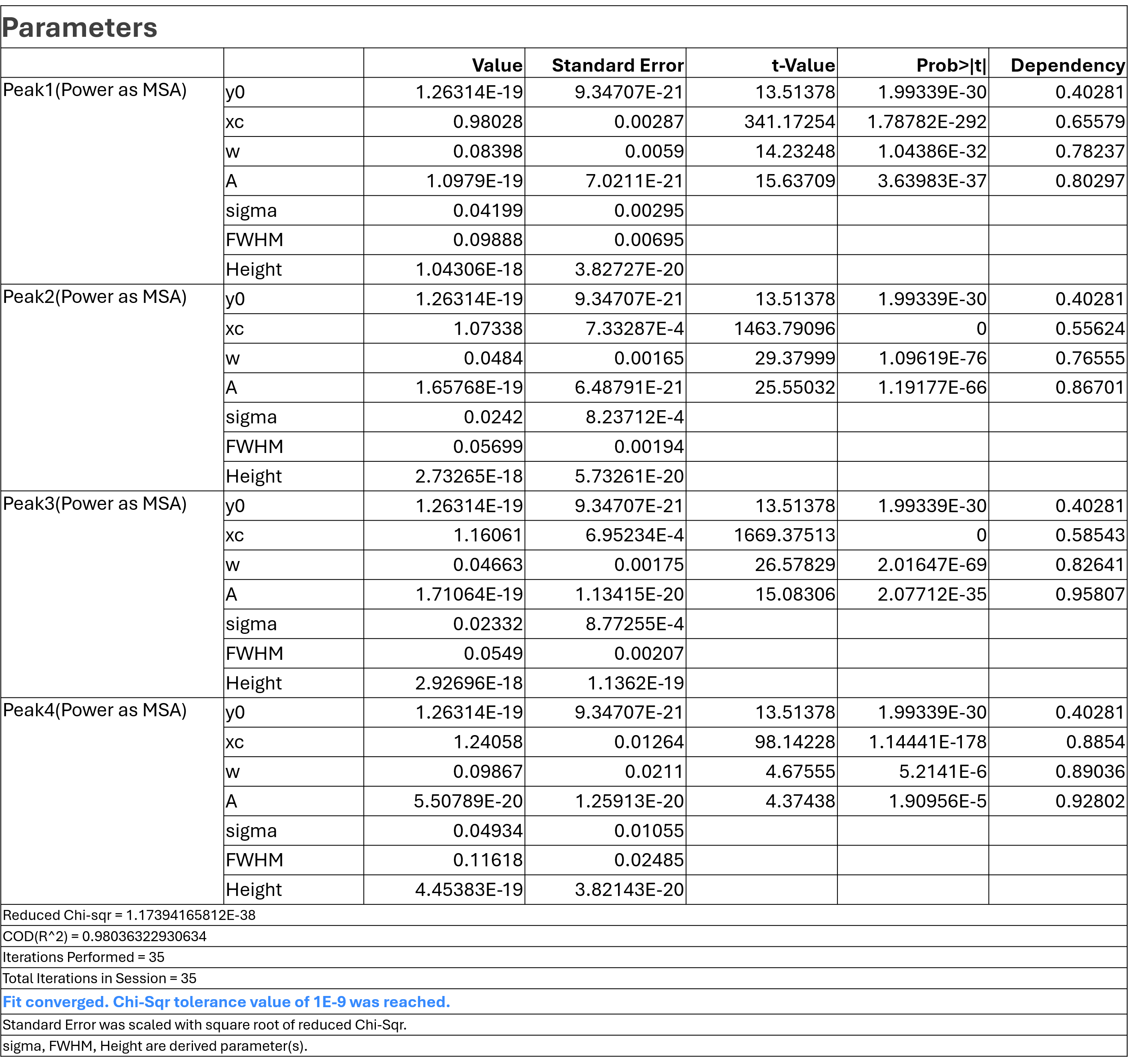
